# T-SCAPE: T-cell Immunogenicity Scoring via Cross-domain Aided Predictive Engine

**DOI:** 10.1101/2025.05.11.653308

**Authors:** Jeonghyeon Kim, Nuri Jung, Jayyoon Lee, Nam-Hyuk Cho, Jinsung Noh, Chaok Seok

**Affiliations:** Department of Chemistry, Seoul National University, Gwanak-gu, Seoul & 08826, Republic of Korea; Graduate School of Data Science, Seoul National University, Gwanak-gu, Seoul & 08826, Republic of Korea; Department of Biomedical Sciences, Seoul National University, Jongro-gu, Seoul & 03080, Republic of Korea; Galux Inc., Gwanak-gu, Seoul & 08738, Republic of Korea

## Abstract

T-cell immunogenicity, the ability of peptide fragments to elicit T-cell responses, is a critical determinant of the safety and efficacy of protein therapeutics and vaccines. While deep learning shows promise for in silico prediction, the scarcity of comprehensive immunogenicity data is a major challenge. We present T-SCAPE, a novel multi-domain deep learning framework that leverages adversarial domain adaptation to integrate diverse immunologically relevant data sources, including MHC presentation, peptide-MHC binding affinity, TCR-pMHC interaction, source organism information, and T-cell activation. Validated through rigorous leakage-controlled benchmarks, T-SCAPE demonstrates exceptional performance in predicting T-cell activation for specific peptide-MHC pairs. Remarkably, it also accurately predicts the ADA-inducing potential of therapeutic antibodies without requiring MHC inputs. This success is attributed to T-SCAPE’s biologically grounded and data-driven multi-domain pretraining. Its consistent and robust performance highlights its potential to advance the development of safer and more effective vaccines and protein therapeutics.

---

T-cell epitope immunogenicity, the ability of short peptide fragments to trigger an immune response by T cells, is a critical factor in adaptive immunity, influencing both protective and pathogenic immune responses (*1, 2, 3*). Understanding the precise immunogenicity of T-cell epitopes has broad implications for various biomedical applications, including the design of effective immunotherapies against cancer, the development of vaccines for infectious diseases (*1, 3, 4*), and strategies to predict and mitigate unwanted immunogenicity in protein-based drugs (*2, 5, 6, 7*).

Traditional experimental methods for identifying T-cell epitopes, such as screening overlapping peptides derived from protein antigens, are costly, time-consuming, and often fail to pinpoint the exact boundaries and immunogenic potency of epitopes (*8, 9*). While *in silico* prediction methods offer a more efficient alternative for T-cell epitope discovery, most current approaches rely on limited data integration, focusing primarily on individual aspects of the T-cell recognition process, such as peptide-MHC binding or T-cell receptor (TCR) and peptide-MHC (pMHC) interaction (*10, 11*). This narrow focus limits both predictive accuracy and generalizability, thereby hindering the development of reliable tools for practical immunogenicity assessment because it neglects critical downstream biological events.

To overcome these limitations, we introduce T-SCAPE, a novel deep learning framework designed to integrate diverse immunogenicity-related data and learn the complex determinants governing T-cell epitope immunogenicity. By employing a modular multi-task learning architecture, T-SCAPE leverages information from multiple domains, including peptide-MHC interactions, TCR-pMHC binding, source organism of peptides, and T-cell activation, to build a more comprehensive and predictive model of T-cell immunogenicity.

## Results

### Model architecture and training strategy

To effectively integrate diverse resources pertinent to T-cell immunogenicity and capture its complex determinants, T-SCAPE employs a modular multi-task learning architecture. This architecture comprises a shared integrative encoder that learns domain-invariant features common across tasks, alongside task-specific encoders that capture complementary, domain-specific information (Fig. 1B, D, F). To promote disentanglement, orthogonality constraints are applied between encoders, encouraging the separation of shared and task-specialized representations. Furthermore, an adversarial loss is employed to guide the integrative encoder toward producing domain-invariant representations by training it to confuse a discriminator tasked with identifying the data source. This mechanism mitigates the risk of overfitting to dataset-specific artifacts—such as peptide length discrepancies between MHC class I and II—and instead promotes learning of fundamental, immunogenicity-relevant features. The model training follows a two-stage process: multi-task pretraining and immunogenicity-specific fine-tuning.

**Figure 1:**
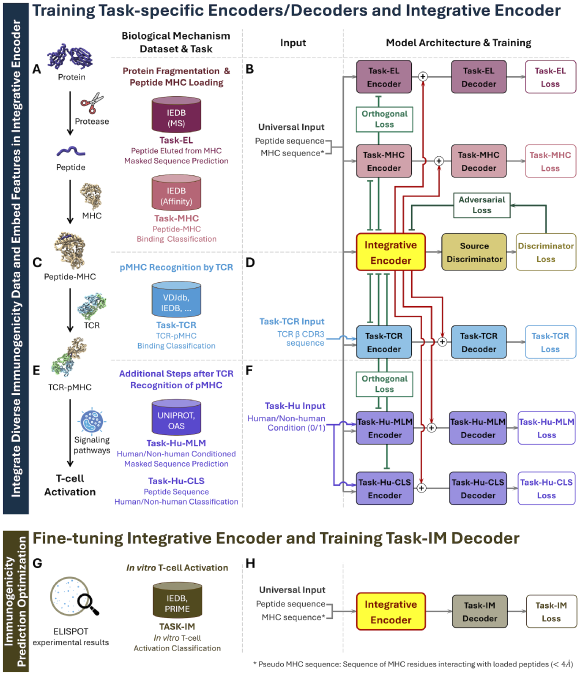
Schematic overview of the T-SCAPE framework architecture and two-phase training strategy. T-SCAPE employs a modular multi-task learning architecture, which features a shared integrative encoder (yellow box) for learning domain-invariant features and complementary taskspecific encoders for extracting domain-specific information. Training is divided into two phases. (**A** to **F**) Pretraining: The integrative and task-specific modules are jointly trained on diverse datasets spanning multiple immunogenicity-relevant tasks, including MHC presentation (Task-EL I, II), peptide–MHC binding affinity (Task-MHC I, II), TCR–pMHC binding (Task-TCR), and peptide “selfness” learning (Task-Hu-CLS/MLM). This phase incorporates task-specific losses from each decoder head, along with orthogonality constraints (light green lines) and an adversarial loss (dark green line) guided by a source discriminator (gold box). This strategy enforces the separation of representations, enhancing generalizability and the learning of fundamental immunogenic features shared across datasets. (**G** to **H**) Fine-tuning: The pre-trained integrative encoder is further optimized with a smaller T-cell activation dataset, using a dedicated Task-IM decoder, to directly predict final immunogenicity.

During the initial multi-task pretraining phase, T-SCAPE is trained simultaneously on diverse immunologically relevant datasets, each aligned with a specific task designed to consolidate broad biological knowledge. Peptide–MHC interaction data, representing the early steps of antigen presentation, are relatively abundant. For example, eluted ligand mass spectrometry (MS) data—such as those from IEDB (*12*), comprising 557,151 class I and 206,055 class II entries and commonly used by models like BigMHC (*11*) and NetMHCpan-4.1 (*10*)—inform “Task-EL (I, II)”, a masked amino acid prediction task modeling peptide processing and MHC presentation (Fig. 1A). In parallel, peptide–MHC binding affinity data—129,973 class I and 516,463 class II measurements, foundational to tools such as MHCflurry-2.0 (*13*)—support “Task-MHC (I, II)”, which predicts binding affinities to capture interaction strength (Fig. 1A). To capture the downstream stages of immune recognition, the critical interaction between TCR and pMHC complexes is addressed through “Task-TCR”, leveraging 65,030 curated data points from IEDB and literature sources (Fig. 1C). This task allows the model to capture the broader process of immune recognition subsequent to peptide-MHC loading, moving beyond the singular focus of interaction-based methods like DLpTCR (*14*). To incorporate concepts of “selfness”, T-SCAPE introduces additional tasks using human-derived protein sequences from the OAS dataset (*15*) and UniProt (*16*). These include “Task-Hu-CLS” (a human vs. non-human classification task, inspired by Hu-mAb (*17*)) and “Task-Hu-MLM” (a species-conditioned masked language modeling task, following ideas from AbNatiV (*18*) and Bio-phi (*19*)) (Fig. 1E).

Following pretraining, the shared integrative encoder is fine-tuned using T-cell activation data to enable final immunogenicity prediction. This data—representing the endpoint of the antigen recognition cascade as measured by functional assays such as ELISPOT—are sourced from repositories including IEDB (*12*), MHC BN (*20*), the NCI database (*21*), and PRIME-2.0 (*22*). Despite its relatively limited size (12,782 data points), this dataset enables direct optimization of the model through “Task-IM” (Fig. 1G). By leveraging rich contextual representations learned during pretraining, T-SCAPE achieves robust immunogenicity prediction—setting it apart from models such as PRIME-2.0 (*23*), which also predict T-cell activation but lack comparable multi-domain pretraining.

### Key architectural features

Our model, T-SCAPE, is built upon a convolutional neural network (CNN) to effectively predict T-cell immunogenicity (Fig. 2A). We chose a CNN-based architecture over transformer or LSTM models because immunogenicity tasks often rely on analyzing short peptide fragments. This characteristic makes the ability of CNNs to capture local sequence features more advantageous than modeling the long-range dependencies required for full-length proteins.

**Figure 2:**
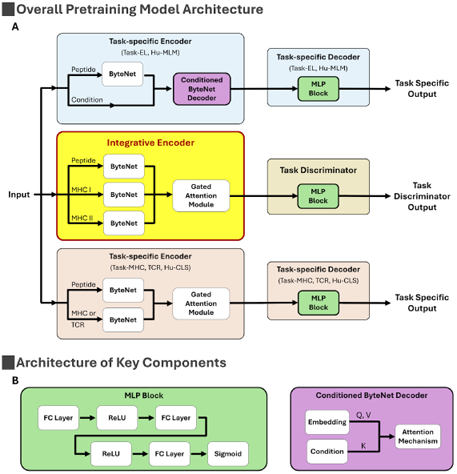
Overview of the T-SCAPE model architecture. **(A)** Overall pretraining model architecture: This diagram illustrates the T-SCAPE’s overall pretraining framework. It highlights the shared integrative encoder and the pathways to various task-specific encoders and decoders. The architecture also includes a task discriminator for adversarial learning, which promotes domain-invariant feature learning. (**B**) Architecture of key components: This panel provides a detailed view of the internal blocks used in the framework. It shows the structure of the Multi-Layer Perceptron (MLP) block used within the decoders and discriminator. It also details a specialized Conditioned ByteNet Decoder block that takes both sequence and a conditioning label as input. For Task-Hu-MLM, this condition is a human vs. non-human label, while for Task-EL, it is MHC information.

The architecture features a shared integrative encoder that processes inputs common to most tasks, along with task-specific encoders for any additional inputs they may require. These encoders are connected by a gated attention module. This mechanism is critical as it allows the model to intelligently focus on the most relevant inputs for a given task while effectively ignoring unnecessary, padded inputs (e.g., ignoring the MHC input for a task that only requires a peptide), ensuring that only the most critical information is utilized.

The resulting feature embeddings from both the shared and task-specific encoders are concatenated and passed to a simple task-specific decoder, which consists of a Multi-Layer Perceptron (MLP) block (Fig. 2B), and generates the final prediction. Separately, the architecture also includes a task discriminator, also an MLP, which takes the output of the integrative encoder. This component facilitates adversarial training, pushing the shared encoder to learn generalized, task-agnostic representations.

### Biological motivation and justification for multi-domain pretraining

To further support the rationale behind our multi-domain training strategy and to gain deeper insights into the relationship between pretraining and fine-tuning datasets, we analyzed label consistency for peptides shared across both stages. Specifically, we identified peptides that appeared in both the pretraining datasets (Task-EL, Task-MHC, and Task-TCR) and the fine-tuning dataset (Task-IM) by detecting exact matches of 9-mer fragments, and then compared their corresponding biological labels (Fig. 3A). This analysis focused strictly on exact 9-mer matches, a stricter criterion than the training/benchmark separation procedure which excluded fragments with fewer than two mismatches. This stricter criterion was used because even a single amino acid substitution can significantly alter immunogenicity outcomes, as demonstrated in Section Case study: Characterizing immunogenic alterations in humanized antibody pairs. Permitting mismatches in this context could obscure the true correspondence between labels and confound interpretation of inter-dataset consistency.

**Figure 3:**
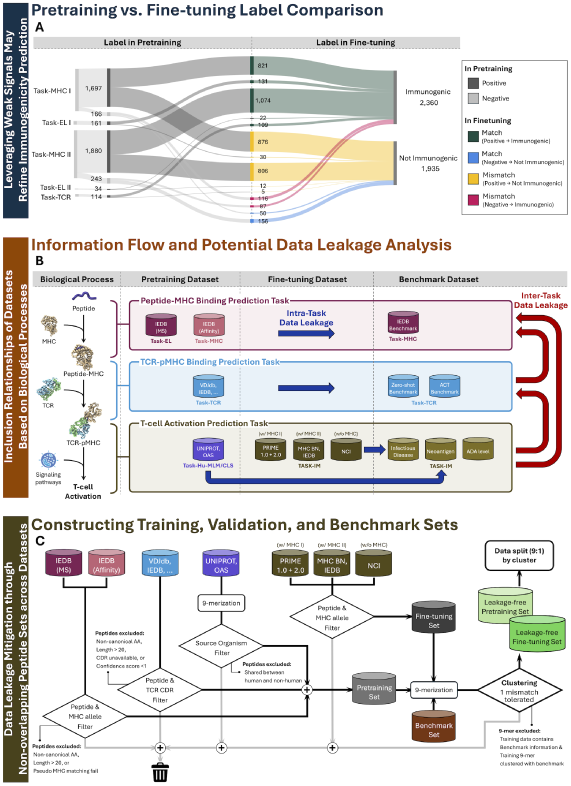
Data-driven motivation and curation strategy for multi-domain learning. This figure outlines the data-centric principles that guided our model development, from initial motivation to rigorous data curation. (**A**) Pretraining vs. fine-tuning label comparison: This Sankey diagram visualizes the concordance between labels from intermediate pretraining tasks and the final immunogenicity labels. The visually distinct mismatched and misaligned flows confirm the complex and non-linear nature of immunogenicity, yet they also reveal a statistically significant underlying relationship (Cramer’s V = 0.0339, p = 0.0265). This finding validates our core hypothesis: to extract the essential information related to immunogenicity from these multi-domain datasets, a precisely-engineered pretraining architecture is necessary. (**B**) Information flow and potential data leakage analysis: This schematic illustrates the hierarchical relationships among the biological processes. Information flows unidirectionally from downstream outcomes, such as T-cell activation, to upstream events like peptide-MHC binding. This is because downstream data inherently encompasses information from all upstream events. This asymmetry guided our stringent data curation to prevent spurious information transfer, thus avoiding inter-task data leakage that would lead to biased model evaluation. (**C**) Dataset curation and splitting workflow: This flowchart details our pipeline for dataset filtering and ensuring leakage-free splits. The process includes tailored preprocessing for each dataset, followed by a stringent 9-mer-based similarity filtering using cd-hit to remove any peptides with fewer than two mismatches between the training and benchmark sets. This rigorous approach ensures a robust and reliable evaluation of the model’s generalizability.

This analysis yielded two key insights that underscore the complexity of T-cell immunogenicity and the necessity of a precisely-engineered multi-domain learning framework.

First, we frequently observed label inversion—cases in which peptides that exhibited strong MHC binding failed to induce significant cytokine release in *in vitro* activation assays. This observation highlights a fundamental limitation of models trained solely on isolated steps of the immunogenicity cascade. It also indicates that the presence of shared peptides across datasets does not inherently result in trivial data leakage; rather, the model must learn to navigate and integrate non-linear, biologically complex relationships to accurately map intermediate signals to final immunogenic outcomes.

Second, while the correlation between intermediate and final labels was relatively weak, it was not negligible. We observed a Cramer’s V of 0.0339 with a statistically significant p-value of 0.0265, indicating a small but meaningful association between the datasets (Section Cramer’s V analysis of label concordance). This finding suggests that even subtle signals from intermediate tasks can provide valuable information for immunogenicity prediction—provided they are integrated effectively within a multi-domain framework.

Taken together, these observations—the complex, non-linear relationship between intermediate processes and final immune activation, along with the weak yet significant correlation observed between them—motivated our specific multi-domain learning strategy. We hypothesized that while these correlations might be too subtle for naive transfer learning, their statistically significant presence provides a valuable signal. Our approach was therefore designed not to rely on strong, direct knowledge transfer, but to strategically exploit these faint inter-task relationships as a form of guided data augmentation. This allows the model to capture and leverage intricate biological dependencies that would otherwise be missed, thereby enhancing the robustness and accuracy of immunogenicity prediction.

### A scrupulous data curation and leakage control strategy

Data leakage is a critical flaw where information from a training dataset unintentionally contaminates the evaluation data, causing a model to appear more accurate than it is on genuinely unseen data. Since data leakage poses a critical challenge in developing and evaluating deep learning models—particularly in multi-domain learning frameworks that integrate data from diverse sources—we implemented a stringent curation and splitting strategy to ensure robust and leakage-free evaluation. To evaluate the risk of data leakage, we first analyzed the biological relationships and the hierarchical structure of information overlap across datasets (Fig. 3B). As expected, the T-cell activation dataset shares content with other activation-related datasets, introducing a risk of intra-task leakage. Moreover, given the sequential nature of the immunogenicity cascade, this dataset inherently incorporates information from upstream processes such as peptide–MHC binding and TCR–pMHC interactions, giving rise to potential inter-task leakage. Similarly, the TCR–pMHC binding dataset subsumes information found in peptide–MHC binding datasets. However, the reverse is not true: peptide–MHC binding datasets do not contain information about downstream processes such as TCR recognition or T-cell activation. Recognizing this asymmetry, we systematically evaluated potential leakage paths from training to benchmark datasets and took explicit steps to eliminate both intra-task and inter-task data leakage.

In line with our stringent strategy to minimize potential data leakage between training and benchmark data across diverse sources, we applied a similarly conservative approach to the human–nonhuman dataset. Although prior studies have suggested that human versus non-human classification provides limited predictive power for immunogenicity (*17, 18, 19*), we treated this dataset as potentially encompassing the entire immunogenicity process, analogous to T-cell activation datasets. This cautious stance was adopted to guard against even subtle forms of leakage, as the dataset may implicitly encode aggregate immunogenicity-related features that transcend individual biological processes.

Prior to applying stringent protocols to mitigate the risks of the intra- and inter-task data leakage, we performed an initial data curation by subjecting each available dataset to a tailored filtering process (Fig. 3C). For the peptide–MHC binding and eluted ligand MS datasets, we excluded peptides containing non-canonical amino acids or exceeding 20 residues in length, in accordance with our model architecture, which accepts inputs up to 20 residues. Shorter peptides were padded to this length to standardize input dimensions. This design accommodates the natural variation in peptide lengths across tasks, particularly the longer peptides commonly found in Task-MHC II. We also removed entries lacking well-defined MHC allele annotations, which are essential for generating accurate pseudo-sequences (*10*). A similar filtering strategy was applied to the TCR–pMHC dataset, excluding peptides based on length or non-standard amino acids. In addition, we discarded samples lacking TCR complementarity-determining region (CDR) sequences or with a VDJdb confidence score below 1 (*24*). For the Human versus Non-human dataset, designed to help the model learn species-specific features, we applied a tailored filtering strategy. Each peptide was segmented into overlapping 9-mer fragments, and the frequency of each fragment was computed separately for human and non-human sequences. Peptides containing any 9-mer that appeared in both species were excluded, ensuring that only peptides uniquely associated with either human or non-human samples were retained. The T-cell activation dataset, used during the fine-tuning stage, was subjected to the same filtering criteria as the peptide–MHC dataset.

Following dataset-specific preprocessing, we applied a stringent filtering protocol to the curated data pool to address the aforementioned intra- and inter-task data leakage. For tasks where a dedicated independent benchmark set was absent, such as for pMHC class II, we first split the data pool into training and benchmark sets and then applied our anti-leakage filters. The filtering process was conducted in the following steps. First, each sample in the training and benchmark sets was decomposed into overlapping 9-mer peptides. Second, we assessed the interand intratask relationship between the training and benchmark set pair to identify potential homology. Subsequently, any training sample was removed if its constituent 9-mer peptides shared sequence homology with any 9-mer peptide in the benchmark set. This homology was defined as having fewer than two mismatches between peptides, a stringent criterion implemented to eliminate even the slightest possibility of data leakage. Notably, this filtering was not performed between Class I and Class II datasets, a decision made to account for their distinct biological pathways. The entire peptide-level filtering procedure was efficiently performed using the CD-HIT clustering algorithm. A more detailed description is provided in Section Sequence identity filtering using CD-HIT.

The 9-mer-based filtering procedure is grounded in the rationale that core T-cell epitopes are often localized within 9-mer regions, a strategy particularly relevant for MHC Class II as it accounts for their relatively fixed binding motif. However, we acknowledge that this approach is less suitable for MHC Class I, which can bind peptides of various lengths with conformations that may bulge from the binding groove. To address the possibility of leakage through these non-consecutive motifs, we conducted a more rigorous analysis for all Class I peptides. Using sequence alignment, we searched for any shared fragments of 8-mers or longer between our training and benchmark datasets. This exhaustive check revealed no such overlaps, confirming the exceptional stringency of our data separation. It is also important to note that our filtering choices do not limit the model’s input length, as it is designed to accept peptide inputs up to 20 amino acids and handles them uniformly via padding.

After constructing a rigorously curated and leakage-free dataset, we split the training data into training and validation subsets using a 9:1 ratio. To ensure independence and prevent information leakage, this split was performed under the same strict anti-leakage protocol as our training-benchmark set separation, which ensured that homologous fragments with fewer than two mismatches were not assigned to different subsets. This deliberate approach simulates a challenging evaluation setting for hyperparameter tuning, thereby enabling a reliable assessment of the model’s true generalization performance. For a detailed breakdown of the data curation process, including dataset composition, filtering statistics, and a comprehensive 9-mer overlap analysis between all training and benchmark sets, please refer to the Supplementary Information (fig. S1).

Our stringent dataset separation strategy, consistently applied across the training, validation, and benchmark splits, surpasses the standards of widely used tools, such as NetMHCpan (*10*) and BigMHC (*11*). While NetMHCpan excludes only exact peptide matches and BigMHC allows overlapping peptide sequences as long as the MHC allele differs, our more conservative approach reflects our critical emphasis on eliminating even subtle forms of data leakage. This ensures robust, unbiased, and generalizable benchmark results.

### T-SCAPE-IM: T-cell epitope immunogenicity prediction for specific peptide-MHC pairs

Understanding the T-cell epitope immunogenicity of peptide–MHC pairs is critical for the identification of neoantigens in cancer immunotherapy and the design of vaccines against infectious diseases. Experimental assays such as ELISPOT are valuable but face significant limitations in high-throughput applications due to the complexities of antigen production (*25*) and the labor-intensive nature of manual spot counting (*26*). In this context, computational prediction models serve as a powerful alternative, enabling substantial reduction in the number of peptide candidates that require experimental validation.

T-SCAPE predicts T-cell epitope immunogenicity by combining an integrative encoder with the Task-IM decoder (Fig. 1H and Fig. 4). This configuration, referred to as T-SCAPE-IM, produces an immunogenicity score from the input peptide and MHC pseudo-sequence, which captures amino acid residues that contact the peptide (*10*). A variant model, t-scape-im, shares the same architecture but is trained solely on the fine-tuning dataset, without leveraging pretraining of the integrative encoder.

**Figure 4:**
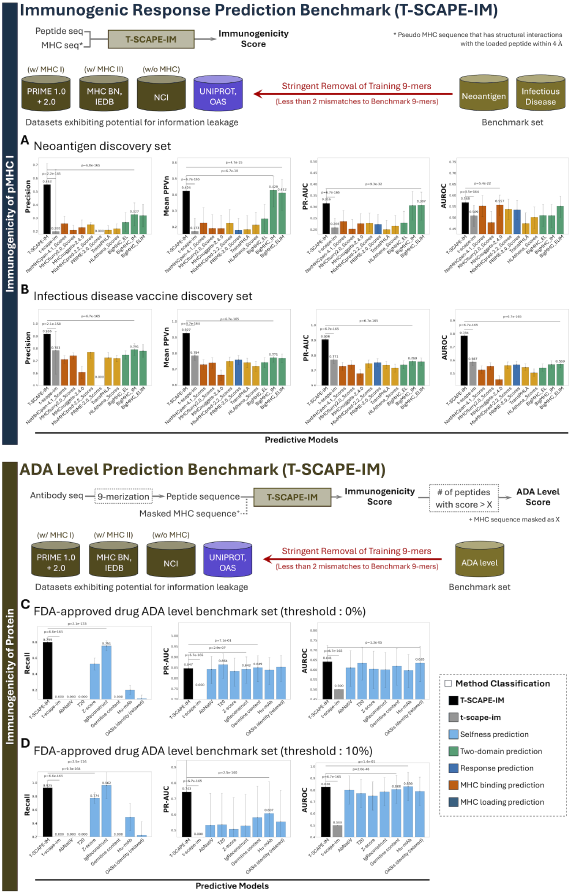
Performance comparison of T-SCAPE-IM on immunogenicity prediction benchmarks. This figure compares the performance of our model, T-SCAPE-IM, against competing methods (color-coded by method type) across two key immunogenicity prediction benchmarks. Panels (**A**) and (**B**) show performance on the neoantigen and infectious disease vaccine discovery sets, respectively, evaluated using Precision, Mean PPVn, PR-AUC, and AUROC. Panels (**C**) and (**D**) evaluate the model’s ability to predict ADA generation levels in therapeutic antibodies under different thresholds. Panel (**C**) uses a strict 0% immunogenicity threshold, where a positive label is assigned if at least one patient develops an ADA response. Panel (**D**) uses a clinically relevant 10% threshold, where a positive label indicates that 10% or more of the patients developed an ADA response. These were assessed using Recall, PR-AUC, and AUROC. A baseline variant, t-scape-im, trained without multi-domain pretraining but sharing the same architecture, is included for comparison to highlight the contribution of our pretraining strategy. Error bars represent 95% confidence intervals from bootstrapping, and statistical significance was assessed using paired Wilcoxon signed-rank tests with Bonferroni correction where applicable.

Model performance was assessed using four key metrics: Mean PPVn (mean positive predictive value among the top n predictions), Precision, PR-AUC (area under the precision–recall curve), and AUROC (area under the receiver operating characteristic curve). The selection of these metrics was guided by their specific relevance to our benchmark setting, which was designed to reflect real-world applications such as neoantigen discovery and vaccine development. First, Precision and PPVn were included because they are crucial for identifying promising candidates for experimental validation in neoantigen discovery and vaccine development. Second, PR-AUC was utilized to verify that high Precision and PPVn values were not obtained at the expense of recall, ensuring the model’s overall robustness. Finally, AUROC was incorporated to maintain consistency with previous studies and facilitate a direct comparison of our model’s performance.

### MHC class I benchmark results on neoantigen and infectious disease vaccine discovery sets

T-SCAPE-IM, along with leading state-of-the-art methods—including BigMHC (*11*), NetMHCpan-4.1 (*10*), MHCflurry-2.0 (*13*), MHCnuggets-2.4.0 (*4*), MixMHCpred-2.2, PRIME-2.0 (*23*), TranspHLA (*27*), and HLAthena (*28*)—was rigorously evaluated on two independent benchmark datasets: (a) a neoantigen discovery set (833 peptides: 178 positives, 655 negatives) and (b) an infectious disease vaccine discovery set (2,093 peptides: 1,488 positives, 605 negatives), as shown in Figure 4A, B.

To ensure that performance evaluations reflect genuine generalization rather than memorization, we applied a strict data leakage prevention strategy that handle inter-task and intra-task data leakage as explained in previous section A scrupulous data curation and leakage control strategy. This rigorous filtering procedure guarantees a robust and fair assessment of model performance. Building on this foundation, we assessed the robustness and statistical significance of each model’s performance through extensive bootstrap analysis and hypothesis testing. For implementation details, see Section Bootstrapping and statistical evaluation in Supplementary Information.

In the neoantigen discovery benchmark (Fig. 4A), T-SCAPE-IM demonstrates significantly and substantially superior performance compared to the non-pretrained variant t-scape-im (*p* < 0.001) across all four metrics, underscoring the critical importance of pretraining. Compared to BigMHC-IM, T-SCAPE-IM showed a small but statistically significant disadvantage in mean PPVn (0.424 vs. 0.429, *p* < 0.001), but demonstrated significantly and substantially superior performance in Precision (0.553 vs. 0.327, *p* < 0.001). Consequently, T-SCAPE-IM also attains a marginally higher PR-AUC, which was statistically significant, compared to BigMHC-ELIM (0.316 vs. 0.307; *p* < 0.001), reflecting its strong Recall performance. Additionally, T-SCAPE-IM exhibits a marginally higher AUROC, which was statistically significant, than MHCnuggets-2.4.0 (0.568 vs. 0.557, *p* < 0.001).

In the infectious disease vaccine benchmark (Fig. 4B), T-SCAPE-IM again demonstrates significantly and substantially superior performance compared to the non-pretrained variant t-scape-im (*p* < 0.001) and establishes its superiority over all existing methods across all metrics. Specifically, it achieved significantly and substantially superior performance compared to BigMHC-IM in both Precision (0.916 vs. 0.791, *p* < 0.001) and mean PPVn (0.927 vs. 0.771, *p* < 0.001). Furthermore, T-SCAPE-IM significantly and substantially outperformed BigMHC-IM in PR-AUC (0.904 vs. 0.760, *p* < 0.001) and BigMHC-ELIM in AUROC (0.784 vs. 0.569, *p* < 0.001).

The error bars in Figure 4, representing 95% confidence intervals computed via bootstrap analysis, reflect the stability of each model’s performance. With the notable exception of the Precision metric in the neoantigen discovery set, T-SCAPE-IM exhibits narrower or comparable error bars relative to other models, indicating robust and consistent performance. This is summarized in Table 1, which presents the confidence interval width for each model and each metric. Notably, compared to the non-pretrained variant t-scape-im, T-SCAPE-IM demonstrates substantially smaller error bars across all metrics, highlighting its enhanced stability in addition to superior predictive accuracy derived from the pretraining.

**Table 1:**
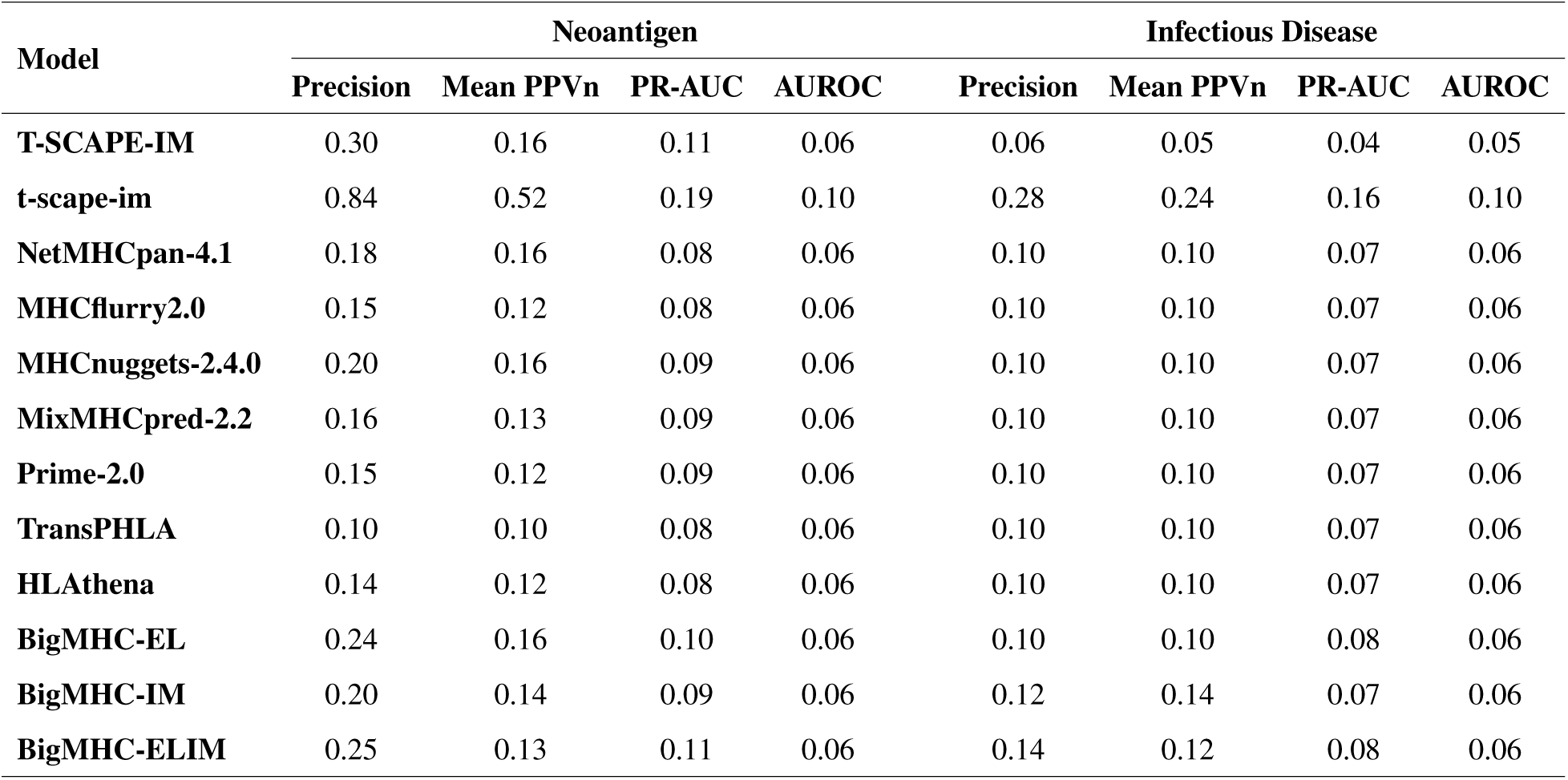
Comparison of predictive stability on Neoantigen and Infectious Disease benchmarks. The values in the table represent the width of the 95% confidence interval. A smaller value indicates more stable prediction performance.

To further dissect the model’s performance, we analyzed its predictions on a per-peptide-length and per-HLA-allele basis. This analysis was conducted on the same independent benchmark sets for neoantigens and infectious diseases from the preceding evaluations. As these benchmarks are specific to MHC class I immunogenicity, the scope of this analysis is naturally confined to peptides 8–11 amino acids in length. For T-SCAPE-IM, bootstrap-based 95% confidence intervals were computed and shaded to indicate the variability in performance.

While the model’s absolute performance metrics varied across different peptide lengths, as shown in Figure 5A, C, T-SCAPE-IM nonetheless consistently outperformed other methods across all tested peptide lengths in both the neoantigen and infectious disease benchmarks. We emphasize that, unlike many existing methods which impose restrictions on peptide length—such as NetMHC-pan (8–14 amino acids) and PRIME-2.0 (8–11 amino acids)—or exhibit performance degradation with increasing peptide length, as observed in BigMHC, our model not only supports peptides up to 20 amino acids in length but also outperforms these existing methods on the benchmark lengths (8–11 amino acids). We attribute this consistent superiority to the pretraining phase, during which the model was exposed to peptides of diverse lengths—an advantage not attainable using the Task-IM fine-tuning dataset alone.

**Figure 5:**
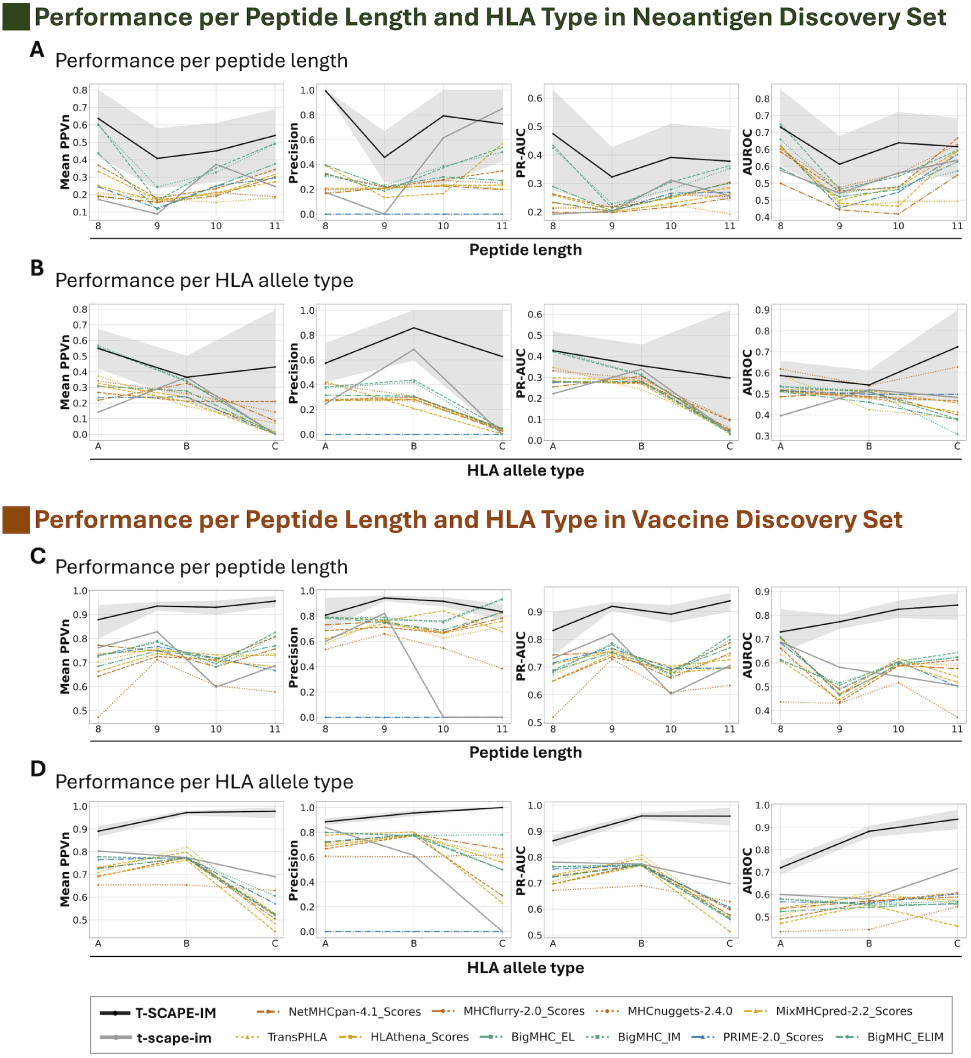
Performance variations by peptide length and HLA type across MHC class I benchmarks. This figure details the performance of T-SCAPE-IM (black line with shaded 95% confidence interval) and competing methods on MHC Class I immunogenicity prediction, stratified by key biological parameters. (**A**) Neoantigen discovery set – performance per peptide length: T-SCAPE-IM exhibits strong performance across all tested peptide lengths (8–11 amino acids) for the neoantigen discovery set, generally outperforming or matching competing methods. (**B**) Neoantigen discovery set – performance per HLA allele type: T-SCAPE-IM maintains competitive performance across diverse HLA allele types (A, B, and C) within the neoantigen discovery set, performing at a level comparable to or exceeding other models. (**C**) Vaccine discovery set – performance per peptide length: Consistent with the neoantigen results, T-SCAPE-IM sustains superior performance across all peptide lengths (8–11 amino acids) for the infectious disease vaccine discovery set, relative to other methods. (**D**) Vaccine discovery set – performance per HLA allele type: T-SCAPE-IM demonstrates exceptional performance across major HLA allele types (A, B, and C) on the infectious disease vaccine discovery set, generally outperforming or matching alternative approaches.

Figure 5B, D illustrates T-SCAPE-IM’s performance across different HLA allele types in both the neoantigen and infectious disease vaccine benchmarks. While the model’s performance was not necessarily robust or stable across different HLA types, it consistently outperformed competing methods in both settings, with its margin of superiority being more pronounced and consistent in the infectious disease benchmark. In the neoantigen benchmark, strong performance is observed for HLA-A, but performance for HLA-B and HLA-C, though superior to other models, is less pronounced. This variation in the performance gap is likely due to disparities in label distribution (table S4).

### MHC class II benchmark results on a rigorously curated, leakage-free benchmark set

In contrast to the preceding benchmark results, which utilized MHC class I datasets, a significant challenge in the MHC class II domain is the absence of a dedicated, independent test set for immunogenicity evaluation. Recognizing this critical gap, we constructed a novel benchmark by rigorously partitioning our internal pMHC class II fine-tuning data. To ensure its integrity and prevent data homology with existing public sets, we first implemented a stringent filtering process, systematically removing any 9-mer with fewer than two mismatches to a 9-mer in any public benchmark (neoantigen discovery, infectious disease vaccine, and ADA sets). The remaining homology-free data was then partitioned into training (80%), validation (10%), and test (10%) sets. This partitioning, guaranteed by our filtering criteria, ensured no sequence overlap between the splits. The final test set comprises a total of 777 samples (436 positive and 341 negative), encompassing a broad range of allele types (13 HLA-DP, 182 HLA-DQ, and 582 HLA-DR).

As shown in Figure 6A, T-SCAPE-IM consistently outperformed existing methods across all metrics on this newly constructed test set. It achieved a Mean PPVn of 0.633, a Precision of 0.611, a PR-AUC of 0.710, and an AUROC of 0.678. This performance represented a statistically significant (*p* < 0.001) improvement over the next-best performing models, including NetMHCIIpan-4.1BA and NetMHCIIpan-4.0EL. Statistical significance was assessed using bootstrap analysis (1,000 resamples) and paired Wilcoxon signed-rank tests with Bonferroni correction.

**Figure 6:**
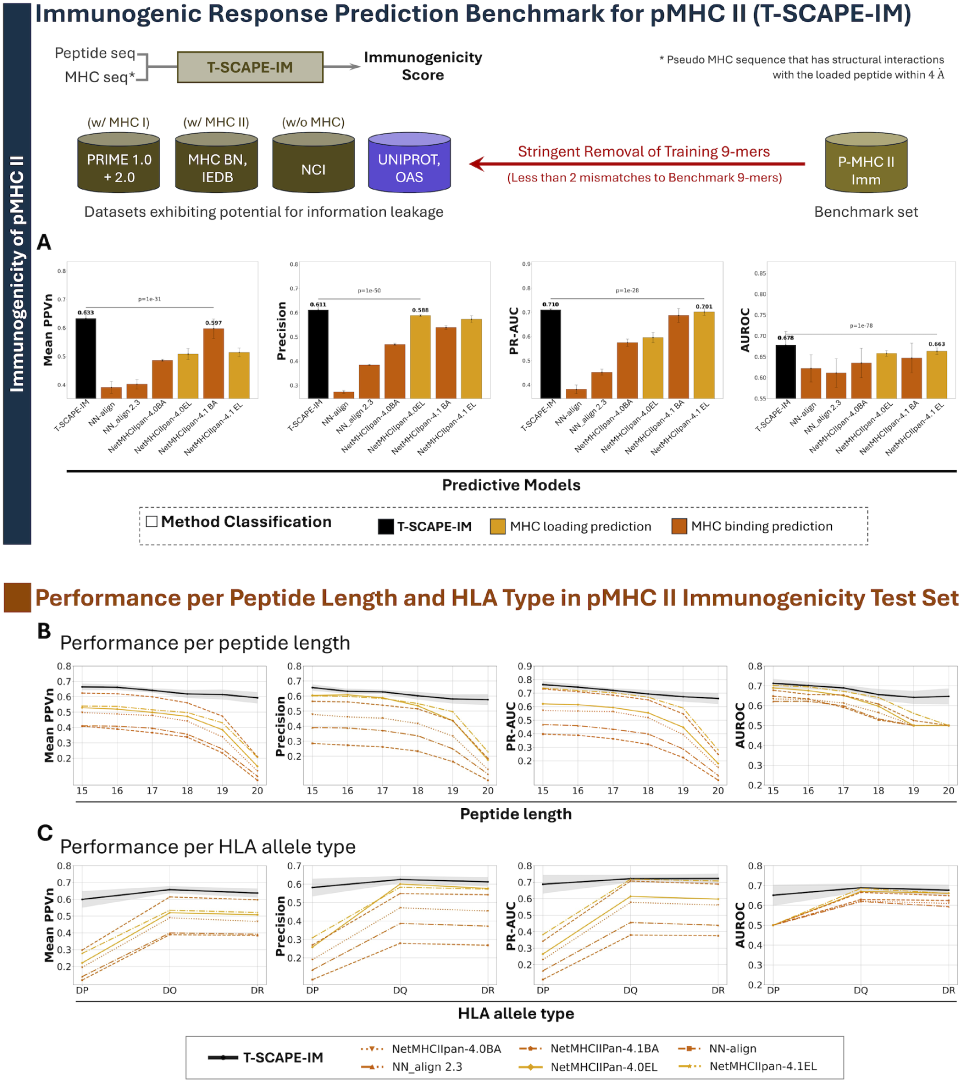
Immunogenicity prediction benchmark for pMHC class II. **(A)** Performance on the pMHC II immunogenicity benchmark set: The top schematic provides a brief overview of the input, model, and output for pMHC II immunogenicity prediction. It also illustrates the data curation process used to construct a rigorous, independent test set. This involved the stringent removal of 9-mers from the training set that had fewer than two mismatches when compared to any 9-mer in the benchmark set, ensuring no homology. The bottom panel shows the performance comparison of T-SCAPE-IM with other MHC class II prediction models on this benchmark, evaluated across Mean PPVn, Precision, PR-AUC, and AUROC. (**B**) Performance per peptide length and (C) performance per HLA allele type: These panels show the model’s performance when stratified by peptide length (15–20 amino acids) and HLA allele type (DP, DQ, and DR), respectively. The shaded areas in both panels represent 95% confidence intervals estimated via bootstrapping.

Furthermore, to specifically evaluate the model’s robustness on longer peptides and diverse alleles of MHC class II, we performed an additional stratified analysis (Fig. 6B, C). This analysis highlighted the model’s ability to process long peptides and diverse MHC class II alleles (HLA-DP, -DQ, and -DR). While most models exhibited a significant performance drop for longer peptides (up to 20 amino acids) where data is sparse, T-SCAPE-IM maintained highly stable and superior performance. Similarly, for the HLA-DP allele, which was the least represented in the dataset and caused performance degradation in other models, T-SCAPE-IM again demonstrated the most robust results, maintaining its predictive power and outperforming all other methods.

### Case study: Modeling immunogenic variation from subtle alterations in pMHC complexes

Figure 7 presents a case study to verify whether T-SCAPE-IM’s predictions align with *in vitro* cell assay results, particularly in instances where minor input variations lead to a complete reversal of outcomes. We found that T-SCAPE’s predictions align sharply with experimental observations of how both single-residue peptide mutations and presenting MHC alleles can dramatically alter immunogenicity. For instance, a single amino acid mutation in a peptide from the SARS-CoV-2 spike glycoprotein results in a sharp contrast in predicted immunogenicity by T-SCAPE (0.9669 vs. 2.95e-05), a finding consistent with experimental validation (Fig. 7A). Similar trends are observed with peptides from Hepatitis D virus and HIV-1 (Fig. 7B, C), highlighting the model’s sensitivity to subtle peptide sequence variations. Beyond peptide mutations, the same peptides from sources such as SARS-CoV2 and Mammarenavirus lassaense also yield drastically different T-SCAPE scores when presented by different MHC alleles, with these scores accurately reflecting their experimentally validated immunogenicity (Fig. 7D to F). These results demonstrate T-SCAPE-IM’s capacity to capture immunogenic variation arising from minimal peptide alterations and HLA context differences.

**Figure 7:**
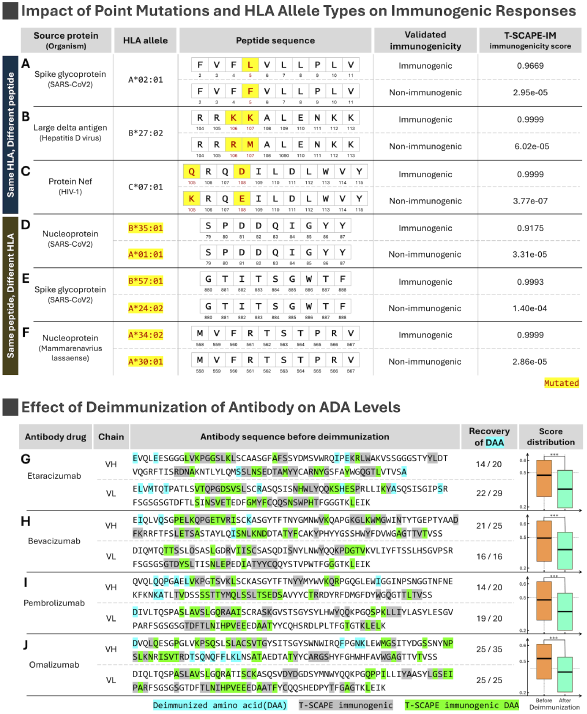
Case studies demonstrating T-SCAPE-IM performance. This figure presents various case studies that highlight T-SCAPE-IM’s predictive capabilities across different immunogenicity scenarios. (**A** to **f**) Impact of point mutations and HLA allele types on immunogenicity: This section illustrates T-SCAPE-IM’s ability to predict changes in immunogenicity due to peptide sequence alterations or HLA variations. (**A** to **C**) Single point mutations: T-SCAPE-IM accurately discriminates the impact of single amino acid point mutations on immunogenicity for antigens such as (**A**) SARS-CoV-2 Spike glycoprotein, (**B**) Hepatitis D virus Large delta antigen, and (**C**) HIV-1 Nef protein. For each case, the predicted T-SCAPE-IM Immunogenicity score aligns with the experimentally validated immunogenicity. (**D** to **F**) HLA allele type variations: The model effectively captures the influence of different HLA allele types on immunogenicity for identical peptides derived from (**D**) SARS-CoV-2 Nucleoprotein, (**E**) SARS-CoV-2 Spike glycoprotein, and (**F**) Lassa virus Nucleoprotein. (**G** to **J**) Effect of deimmunization on predicted ADA levels: This section demonstrates T-SCAPE-IM’s utility in assessing therapeutic antibody deimmunization. For each antibody drug—(**G**) Etaracizumab, (**H**) Bevacizumab, (**I**) Pembrolizumab, and (**J**) Omalizumab—the model identifies experimentally mutated residues (highlighted in the sequence) and shows a clear reduction in predicted ADA-level scores between the sequences before and after deimmunization. Specifically, these ADA-level scores represent the distribution of immunogenicity scores derived from applying a 9-mer peptide window across the entire antibody sequence. Statistically significant score differences (*p* < 0.001) are denoted by ***. Amino acid color codes and residue numbering conventions are provided within the figure panels.

### T-SCAPE-IM: Application to anti-drug antibody level prediction for therapeutic antibodies

Minimizing the immunogenicity of therapeutic antibodies, as measured by anti-drug antibody (ADA) production, is crucial for ensuring drug safety and efficacy. While selfness-based methods have been commonly used to estimate ADA risk, T-SCAPE-IM offers a distinct approach by classifying an antibody’s immunogenicity through a two-step process based on immunogenicity scores. First, we classify a peptide as immunogenic if its score exceeded the 0.5 threshold, which represents the standard boundary for a binary classifier trained with Binary Cross-Entropy loss. Second, an antibody is deemed immunogenic if more than 20% of its constituent peptide fragments are classified as immunogenic, a criterion selected based on established practices in a previous study (*19*). To ensure methodological reliability and reproducibility, we adopted the score threshold and immunogenic peptide ratio from an established study; however, we anticipate that tuning these values on an appropriate dataset could further enhance the model’s performance.

Given the critical importance of identifying all potential immunogenic candidates early in this process, we emphasize Recall as the primary evaluation metric. It directly reflects the model’s ability to correctly identify all potential immunogenic antibodies. To account for the trade-off between high Recall and Precision, we also calculated PR-AUC to evaluate the overall performance. Finally, AUROC was included as a standard metric to facilitate a fair performance comparison with other studies.

### Benchmark results on ADA-level benchmark set

We evaluated the predictive performance of T-SCAPE-IM, its non-pretrained baseline (t-scape-im), and several established selfness-based methods—including AbNatiV (*18*), T20 (*29*), Z-score (*30*), IgReconstruct (*31*), AbLSTM (*32*), Germline content (*33*), Hu-mAb (*17*), MG Score (*34*), and OAsis (*19*)—on the ADA-level benchmark dataset of FDA-approved antibody drugs (size: 216; Threshold 0%: positive - 169, negative - 47; Threshold 10% : positive - 53, negative - 163) (*17, 18, 19*).

The benchmark dataset was evaluated using two distinct thresholds (0% and 10%) to reflect different clinical priorities. The 0% threshold represents a stringent, highly conservative standard for identifying any potential anti-drug antibody (ADA) risk in a single individual, prioritizing maximum drug safety. In contrast, the 10% threshold provides a more clinically relevant and affordable benchmark, representing a level of ADA generation that may be deemed acceptable for certain therapeutic applications.

To maintain data integrity and prevent leakage, we applied a strict data leakage prevention strategy that handle inter-task and intra-task data leakage as explained in previous section A scrupulous data curation and leakage control strategy. This meticulous filtering procedure guarantees a robust and fair assessment of model performance. To assess the statistical significance and stability of our results on this rigorously filtered dataset, we conducted extensive bootstrap analysis and hypothesis testing. For implementation details, see Section Bootstrapping and statistical evaluation in Supplementary Information. Overall, the analysis confirmed that T-SCAPE-IM consistently and significantly outperformed all other evaluated methods across most of the evaluated metrics.

In the 0% immunogenicity threshold setting, as shown in Figure 4C, the non-pretrained baseline, t-scape-im, failed to identify any immunogenic antibodies, classifying all sequences as non-immunogenic. Consequently, it achieved Recall and PR-AUC scores of 0 and an AUROC of 0.5, equivalent to random guessing. In stark contrast, T-SCAPE-IM achieved a marginally higher Recall, which was statistically significant, (0.799) compared to all other models, including the next-best, IgReconstruct (0.751, *p* < 0.001). While achieving this high Recall, T-SCAPE-IM maintained a strong balance between Precision and Recall, with a PR-AUC of 0.847 that was only marginally lower, a statistically significant difference, than T20 (0.864, *p* < 0.001). Its AUROC of 0.641 was also marginally higher, a statistically significant difference, than OASis identity’s (0.635, *p* < 0.001).

Furthermore, to evaluate performance under a standard with greater clinical relevance, we reanalyzed the data using a 10% immunogenicity threshold (Fig.4D). This threshold significantly reduces the number of positive samples, creating a more challenging classification task. Under this condition, while the baseline variant continued to exhibit poor performance, T-SCAPE-IM demonstrated a remarkably well-balanced and robust profile. Although its Recall (0.925) was marginally lower than that of IgReconstruct (0.962), it achieved a substantially higher PR-AUC of 0.743 compared to all other models, including the next-best Hu-mAb (0.607). This superior PR-AUC is particularly noteworthy, as it suggests T-SCAPE-IM has a significant advantage in precision. Rather than indiscriminately flagging sequences to inflate recall, the model accurately identifies true positives, showcasing its reliable classification ability in a low-positive-rate setting. Lastly, its AUROC of 0.826 remained highly competitive and was only marginally lower than that of Hu-mAb (0.830), confirming its robust performance in this clinically relevant scenario.

The 95% confidence intervals of the performance metrics indicate the stability and robustness of each model. T-SCAPE-IM consistently exhibits narrower or comparable error bars relative to other models across both the 0% and 10% threshold settings, indicating robust and stable performance, as shown in Table 2, which details the confidence interval width for each model and metric. This stability of T-SCAPE-IM stands in stark contrast to the non-pretrained variant, t-scape-im. While the non-pretrained model exhibited low variation in its performance metrics, this apparent stability was a direct result of its consistent failure to produce meaningful predictions. This highlights that T-SCAPE-IM’s enhanced stability, in addition to its superior predictive accuracy, is a direct result of its multi-domain pretraining.

**Table 2:** Comparison of predictive stability on the ADA benchmark. The table shows the width of the 95% confidence interval for each model’s performance on this dataset at two different t<u>hresholds. A smaller value indicates more stable prediction performance.</u>

| Model | Threshold 0% |  |  | Threshold 10% |  |  |
| --- | --- | --- | --- | --- | --- | --- |
|  | Recall | PR-AUC | AUROC | Recall | PR-AUC | AUROC |
| **T-SCAPE-IM** | 0.22 | 0.08 | 0.13 | 0.11 | 0.21 | 0.11 |
| **t-scape-im** | 0.00 | 0.00 | 0.00 | 0.00 | 0.04 | 0.00 |
| **AbNatiV** | 0.24 | 0.09 | 0.14 | 0.18 | 0.20 | 0.12 |
| **T20** | 0.25 | 0.08 | 0.14 | 0.20 | 0.20 | 0.12 |
| **Z-score** | 0.30 | 0.10 | 0.15 | 0.08 | 0.20 | 0.12 |
| **IgReconstruct** | 0.24 | 0.09 | 0.13 | 0.20 | 0.20 | 0.12 |
| **Germline content** | 0.28 | 0.08 | 0.14 | 0.20 | 0.20 | 0.12 |
| **Hu-mAb** | 0.26 | 0.10 | 0.15 | 0.20 | 0.20 | 0.12 |
| **OASis identity (relaxed)** | 0.26 | 0.10 | 0.14 | 0.20 | 0.20 | 0.12 |

### Case study: Characterizing immunogenic alterations in humanized antibody pairs

To evaluate whether T-SCAPE-IM can effectively distinguish between pre-humanization antibody sequences and their post-humanization counterparts developed through CDR grafting, we analyzed representative cases from (*19*). This analysis serves as a case study to demonstrate the model’s performance in a practical, real-world scenario, contrasting with our preceding benchmark evaluations that enforced a strict zero-overlap policy. The pre-humanization were sourced from the ADA benchmark dataset, ensuring that they did not contain any 9-mers present in our training data. While their corresponding humanized counterparts exhibit a minimal overlap with our training set, and an especially negligible overlap with the fine-tuning set, we argue this does not compromise the validity of the findings for a practical application. For a detailed breakdown of this overlap, please refer to table S1.

Our method for identifying key immunogenic regions at the residue level is as follows. To translate the 9-mer predictions to residue-level highlights, a residue was marked as immunogenic (green) if it was part of at least two 9-mer regions with a score exceeding 0.5. Residues marked “T-SCAPE Immunogenic” are those predicted as immunogenic by our model using the pre-humanized sequence as input, while “T-SCAPE Immunogenic DAA” are the subset of those predictions that align with experimentally confirmed deimmunizing amino acid (DAA) substitution sites (shown in cyan). Since this DAA recovery analysis was performed exclusively on pre-humanization sequences, which have no overlap with the training data, there is no concern of performance inflation. Using this approach, T-SCAPE-IM successfully recovers 70–100% of DAAs in Etanercept, Bevacizumab, Pembrolizumab, and Omalizumab.

To further validate our model, we compared the overall immunogenicity profiles of the sequences. The score distribution for each sequence is composed of immunogenicity scores for all possible 9-mers generated via a sliding window; the total number of 9-mers (N) is specified in the table S1. We analyzed these distributions by comparing the pre-humanization sequences against their post-humanization counterparts (Fig. 7G–J). A two-sided t-test confirmed that the distributions were statistically distinct. Despite a minimal overlap with the training data (and a negligible one with the fine-tuning set), the post-humanization sequences used for this analysis are not expected to substantially inflate performance metrics. Critically, the pre-humanization sequences have zero overlap, ensuring a valid baseline. The distribution of immunogenicity scores before and after deimmunization clearly distinguishes immunogenic from non-immunogenic variants, validating T-SCAPE-IM’s ability to capture critical immunogenicity-related signals.

### T-SCAPE-IM: Dataset ablation study

To dissect the contribution of each pretraining data source and its corresponding task to T-SCAPE’s predictive capabilities, we performed a comprehensive ablation study. For each ablated variant, we systematically excluded a specific dataset and its associated task during the pretraining phase. Concurrently, the task-specific encoder and decoder corresponding to the ablated task were removed from the model architecture, while the shared integrative encoder and the remaining task-specific modules were retained. Following this modified pretraining, all ablated models underwent the identical fine-tuning procedure as the full T-SCAPE-IM model, utilizing the integrative encoder and the Task-IM decoder on the same fine-tuning dataset.

This ablation process yielded four key variants: T-SCAPE-IM (∼Hu) (excluding human-derived protein data and Task-Hu), T-SCAPE-IM (∼MHC I) (excluding MHC class I binding and eluted ligand MS datasets and related tasks), T-SCAPE-IM (∼MHC II) (excluding MHC class II binding and eluted ligand MS datasets and related tasks), and T-SCAPE-IM (∼TCR) (excluding TCR–pMHC binding data and Task-TCR).

The impact of these ablations was evaluated across three distinct settings (Fig. 8): (a) the neoantigen discovery set (MHC class I-associated), (b) the infectious disease vaccine discovery set (MHC class I-associated), and (c) the FDA-approved drug ADA-level benchmark set (closely linked to MHC class II immunogenicity). We note that (a) the neoantigen discovery dataset and (b) the infectious disease vaccine discovery dataset consist of experimental data measuring CD8^+^ T-cell activation, which is associated with MHC class I pathways. In contrast, (c) the FDA-approved drug ADA-level benchmark dataset captures the prevalence of anti-drug antibody induction in patients administered with therapeutic antibodies. Since antibody production is driven by B-cell activation, which is in turn supported by CD4^+^ T-cell activation, this dataset is primarily associated with MHC class II pathways. For each model variant, we visualized performance using the five metrics reported earlier (Mean PPVn, Precision, PR-AUC, AUROC, and Recall), incorporating error bars representing 95% confidence intervals (derived from bootstrapping) and p-value annotations (paired Wilcoxon signed-rank test with Bonferroni correction) to assess the statistical significance of performance differences relative to the full T-SCAPE-IM model.

**Figure 8:**
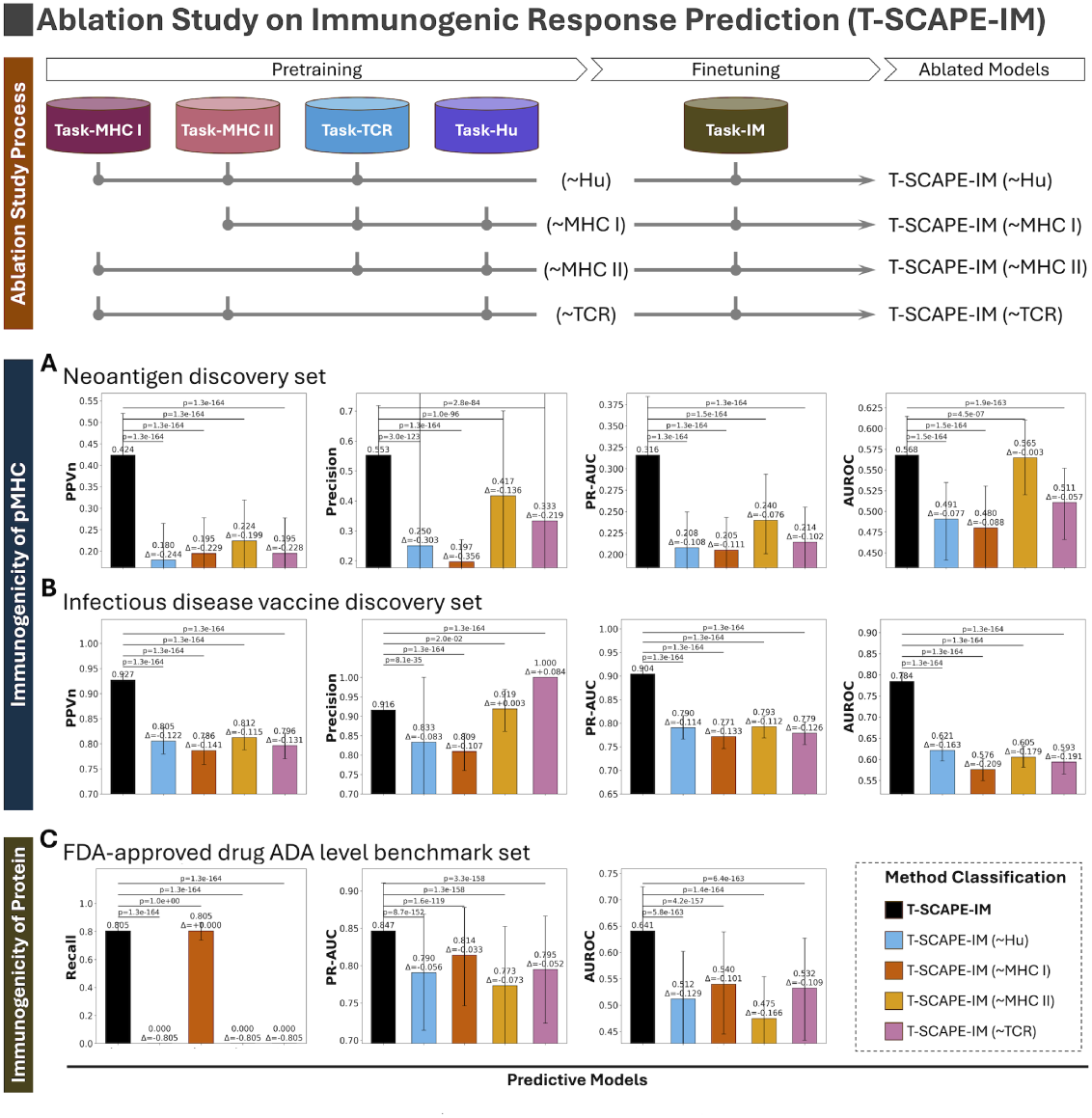
Ablation study on pretraining component contributions. This figure presents an ablation study to evaluate the contribution of each pretraining task to T-SCAPE-IM’s final performance. Ablated variants of the model were generated by excluding a specific pretraining task (∼Hu, ∼MHC I, ∼MHC II, ∼TCR), while all other components—including the architecture and fine-tuning process—remained identical to the full model. The performance of these variants is compared on three key benchmarks: (**A**) the neoantigen discovery set, (**B**) the infectious disease vaccine discovery set, and (C) the ADA-level benchmark set. All results include 95% confidence intervals (estimated via bootstrapping) and are annotated with p-values from a paired Wilcoxon signed-rank test with Bonferroni correction to indicate statistical significance.

The ablation study results confirm the meaningful contribution of all pretraining tasks to T-SCAPE’s overall performance, albeit with varying degrees of influence depending on the specific downstream prediction task. Notably, all comparisons between the full model and the ablated variants revealed statistically significant performance reductions (*p* < 0.001 after Bonferroni correction).

Among the pretraining tasks, Task-Hu exhibited the broadest impact across all benchmarks, likely attributable to the high sequence diversity of the human protein data and its weak yet informative signals related to self/non-self discrimination, a crucial aspect of immunogenicity. This ablation study provided further evidence that the model effectively learns biologically relevant information, showing that removing a task relevant to a specific MHC class caused the most significant performance degradation in its corresponding benchmark set. Specifically, the ablation of Task-MHC I led to the most substantial drops in the CD8^+^ T-cell epitope prediction tasks (Fig. 8A, B), while the removal of Task-MHC II had the most pronounced impact on the CD4^+^ task (Fig. 8C). Interestingly, we also observed a significant performance reduction in CD8^+^ tasks even when Task-MHC II were ablated. Conversely, the CD4^+^ task also showed a significant performance drop when Task-MHC I were removed. We believe this is primarily due to the loss of a data augmentation effect between the Class I and Class II peptide-MHC interaction tasks. Although their peptide-binding grooves are different, the problem is fundamentally one of molecular binding, suggesting that each task provides meaningful information to the other that is crucial for robust performance. While Task-TCR contributed to overall performance, its ablation resulted in a relatively small performance decline. This suggests a limitation in the model’s current architecture, which may not be fully optimized to capture the high complexity and variability of the TCR CDR regions.

## Discussion

In this study, we introduced T-SCAPE, a comprehensive deep learning framework designed to predict T-cell epitope immunogenicity by integrating diverse biological information. T-SCAPE demonstrated exceptional performance via its fine-tuned module (T-SCAPE-IM) across its primary immunogenicity prediction tasks.

### Performance insights and key strengths

T-SCAPE achieved state-of-the-art results in predicting immunogenicity for specific peptide-MHC pairs and delivered top-tier performance in assessing therapeutic antibody immunogenicity. This competitive performance across both domains is a testament to our multi-domain pretraining scheme, which spans broader biological mechanisms than existing methods. This holistic approach acts as an effective data augmentation, enhancing the model’s ability to distinguish between immunogenic and non-immunogenic sequences. By learning the inter-dependencies of the full biological process, T-SCAPE recognizes patterns viable across the entire pathway, making it less prone to false positives.

For T-cell activation, the model demonstrated enhanced performance in neoantigen identification, particularly with significant improvements in overall classification accuracy. We attribute this to our multi-domain pretraining, which integrates signals across the entire immunogenicity cascade. Notably, the performance margin on the infectious disease benchmark was even greater, suggesting the model is particularly well-suited for identifying pathogen-derived epitopes. This strong performance is attributed to the incorporation of the human-nonhuman discrimination domain during pretraining, which enables the model to internalize features that distinguish self from non-self at the sequence level, thereby enhancing its ability to identify epitopes from foreign pathogens.

The model also delivered competitive predictive performance in assessing therapeutic antibody immunogenicity, as measured by anti-drug antibody production. Importantly, this was achieved without the need for MHC pseudo-sequences, a key advantage attributed to the robust pretraining on diverse, multi-domain datasets, including the MHC-independent Task-Hu. These findings underscore T-SCAPE’s significant potential for accurate prediction in the crucial domain of therapeutic antibody development.

### Architectural considerations and model enhancements

Our use of an orthogonality constraint is motivated by the need to effectively separate shared and task-specific information within our multi-task framework. This approach is known to be highly efficient at extracting common representations from a diverse set of tasks, even those without strong explicit commonalities (*35*). By enforcing orthogonality, we encourage the shared encoder to learn features common to all tasks, while the task-specific encoders are guided to focus only on distinct, non-overlapping features. The degree to which this conceptual separation is achieved is quantified by the orthogonal loss curve, which demonstrates a successful reduction in representational overlap during pretraining (fig. S7B).

We incorporated a species-conditioned masked language modeling (Task-Hu-MLM) task to teach the model a more nuanced understanding of human versus non-human sequences. In this task, the model is prompted to predict masked amino acids in a sequence while being provided with a conditioning label indicating whether the sequence is of human or non-human origin (human - 1, other species - 0). Our motivation for this design stems from the observation that while many models use species information, they often do so through a simple classification task (e.g., predicting the species of an entire sequence) (*17, 19*). We hypothesized that conditioning a generative task like masked language modeling with the species label would provide a more powerful data augmentation effect. This approach compels the model to learn not just the high-level features of a species, but also the specific, contextual amino acid patterns that define it, allowing for a richer and more detailed representation of species-specificity.

While our current architecture has several strengths, a key area for future improvement lies in the TCR module. Our model currently relies solely on the CDR3 β sequence—a simplification that, while common, has been shown to be suboptimal (*36*). Although the model’s encoder architecture is flexible enough to be adapted to process the CDR3 α chain as well, the scarcity of largescale, paired-chain TCR data precluded this addition. This design choice thus aligns with our goal of treating the TCR task as auxiliary. Future iterations, contingent on the availability of more comprehensive datasets, could incorporate both the CDR3 α and β chains, along with other CDR loops, to create a more complete TCR representation. This would likely boost performance on TCR-centric prediction tasks.

### Practical applications, limitations, and future directions

Regarding the model’s real-world readiness, we believe T-SCAPE is suitable for immediate use as an auxiliary tool in drug candidate prioritization or de-immunization workflows. For applications such as vaccine design or identifying a promising candidate sequence, the model’s ranking scores can be used to screen and prioritize sequences for experimental validation. In this context, metrics like Precision and Mean PPVn suggest the potential success rate of the top-ranked candidates. Conversely, for de-immunization tasks where the goal is to flag any potentially problematic sequences, Recall serves as a critical metric. In both scenarios, the model can significantly reduce the experimental search space, saving time and resources.

However, for this model to become a predictive tool that can be used independently without separate experimental verification, further improvements are necessary. The primary limitations are data-related; future performance gains would be most significantly advanced by: (1) the availability of more ELISPOT data covering a broader diversity of HLA alleles, (2) larger and more comprehensive clinical datasets, such as ADA datasets, and (3) new datasets for other biological steps in the immunogenicity cascade. In addition to these data requirements, we anticipate that leveraging extensive computational resources to perform exhaustive hyperparameter optimization for pretraining will further enhance the model’s performance. Despite these limitations, we anticipate T-SCAPE’s application will accelerate key processes in biomedical discovery, ultimately contributing to the development of safer and more effective treatments.

### T-SCAPE: A holistic approach to immunogenicity prediction

In this study, we introduced T-SCAPE, a comprehensive deep learning framework designed to predict T-cell epitope immunogenicity by integrating diverse biological information. The broad effectiveness of T-SCAPE across various benchmark schemes underscores the power of our pretraining strategy. This strategy, which integrates a curated array of datasets through a modular multi-task architecture, allows the model to extract crucial information related to immunogenicity from the complex causal relationships among different biological domains. This novel approach enables the model to make robust and holistic predictions. In doing so, it outperformed existing methods across multiple benchmark schemes and various metrics by learning the inter-dependencies across the entire immunogenicity cascade. While acknowledging the critical need for more extensive and diverse biological data to unlock its full potential, the T-SCAPE framework provides a powerful foundation for future research. The public availability of T-SCAPE via a web server (https://galaxy.seoklab.org/design/t-scape) is designed to facilitate broad accessibility and accelerate its application in real-world biomedical discovery, ultimately paving the way for the development of safer and more effective treatments.

## Materials and Methods

### Dataset and task

We compiled all publicly available data, as detailed in Table S2, S3. For the EL task, we collected MHC eluted ligand MS data from IEDB (*12*) using the following filtering criteria: “Epitope Structure: Linear Sequence”, “Include Positive Assays”, “No T cell assays”, “No B cell assays”, “Host: Homo Sapiens (human)”, and “MHC Assays: MHC ligand elution assay”. Then the dataset underwent prefiltering and removal of any overlapping sequences with the benchmark set, as described in Section A scrupulous data curation and leakage control strategy. We note that we did not allow any overlapping with the benchmark set in either the training set or validation set by using the stringent filtering method outlined in Section Sequence identity filtering using CD-HIT. This resulted in 557,151 data points for HLA class I and 206,055 for HLA class II. As all entries were positive, we employed a Masked Language Modeling task by masking the peptide and conditioning on the MHC pseudo sequences (*37*). The dataset was divided into training and validation sets in a 9:1 ratio, ensuring that peptides in the validation set did not overlap with those in the training set using the stringent filtering described in Section Sequence identity filtering using CD-HIT.

For the MHC task, we utilized IEDB (*12*) to obtain the peptide-MHC binding affinity dataset with similar filtering criteria: “Epitope Structure: Linear Sequence”, “Include Positive Assays”, “Include Negative Assays”, “No T cell assays”, “No B cell assays”, “Host: Homo Sapiens (human)”, and “MHC Assays: MHC binding assay”. Then the dataset underwent prefiltering and removal of overlapping sequences with the benchmark set as described in Section A scrupulous data curation and leakage control strategy. We note that we did not allow any overlapping with the benchmark set in either the training or validation set by applying the stringent filtering method as described in Section Sequence identity filtering using CD-HIT. This process yielded 129,973 data points for HLA class I and 516,463 for HLA class II. Data were labeled based on the “Assay - Qualitative Measurement” column: entries containing “positive” were labeled as positive, and those containing “negative” were labeled as negative. This resulted in 57,976 positive and 71,997 negative data points for HLA class I, and 66,424 positive and 450,039 negative data points for HLA class II. We framed the task as a simple binary classification problem, using peptide and MHC sequences as input. The dataset was split into training and validation sets in a 9:1 ratio, ensuring no overlap between peptides in the training and validation sets using the stringent filtering described in Section Sequence identity filtering using CD-HIT.

For the TCR sequence, we utilized positive sets gathered from (*38*), which compiled information from various public data sources, including VDJdb (*24*), IEDB (*12*), McPAS-TCR (*39*), ImmuneCODE (*40*), TBAdb (*41*), and 10X Genomics (*42*). Preprocessing followed the exact methods described by (*38*). The negative set for training was gathered from (*43*), which includes TCR CDR3β sequences derived from healthy donors, a common method to create negative sequences (*44*). From this, we selected CDR3 β sequences with 20 amino acids in length to align with the positive dataset’s sequence length of 20 amino acids. Non-functional CDR3 β sequences were randomly sampled and shuffled with epitopes from the positive sequences to create a balanced 1:1 ratio of positive to negative classes. We note that only CDR3β sequences are utilized in our study, as CDR3α sequences are difficult to obtain. Moreover, several existing *in silico* methods for predicting TCR–peptide–MHC interactions also rely solely on CDR3β sequences (*44, 14*). Then the dataset underwent prefiltering and removal of any overlap with the benchmark set as described in Section A scrupulous data curation and leakage control strategy. We note that we did not allow any overlapping with the benchmark set in either the training or validation set by applying the stringent filtering method described in Section Sequence identity filtering using CD-HIT. This process resulted in 65,030 positive and 65,030 negative data points, totaling 130,060 sequences. We framed this as a binary classification task using peptide and TCR CDR3β sequences as input. The dataset was split into training and validation sets in a 9:1 ratio, ensuring no overlap between peptides in the training and validation sets using the stringent filtering described in Section Sequence identity filtering using CD-HIT.

For the Hu-MLM and Hu-CLS tasks, datasets of human-originated and non-human-originated protein sequences were obtained from UNIPROT (*16*) and OAS (*15*). The sequences were segmented into 9-mer peptides using a sliding 9-mer window applied across each protein sequence. The frequency of each peptide in human and non-human proteins was then counted and the fraction was calculated as in Equation 1. To train the model to distinguish between human and non-human peptides, we selected peptides that i) appeared more than five times and ii) had a fraction value of exactly 0 (entirely non-human) or 1 (entirely human). We adopted this criterion as peptides with fewer than five occurrences may not reliably represent species specificity, and peptides found in both human and non-human sequences could introduce ambiguity in training.

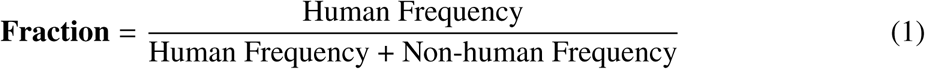

For UNIPROT, given that 400,000 peptides had a fraction of 1, we balanced the dataset by including an equivalent number of non-human peptides. For OAS, we applied the same balancing strategy, resulting in sets of 40,000,000 peptides with a fraction of 1 and 40,000,000 peptides with a fraction of 0. Due to the large size of the dataset, we reduced it to 4,000,000 peptides per class for computational efficiency. We performed a Masked Language Modeling (MLM) task, masking peptides and predicting the masked regions conditioned on whether the sequence was from human or not. Additionally, we conducted a binary classification to distinguish whether the input sequence was human or not. Then the dataset underwent prefiltering and removal of any overlapping sequences with the benchmark set as described in Section A scrupulous data curation and leakage control strategy. We note that we did not allow any overlapping with the benchmark set in either the training or validation set, applying the stringent filtering method as detailed in Section Sequence identity filtering using CD-HIT. The dataset was split into training and validation sets in a 9:1 ratio. To create a hard split validation set, peptides were ordered by frequency, with the last 10% designated as the validation set and the top 90% as the training set. This ensured that the validation set presented a greater challenge compared to the training set.

For the task IM-I, a cytokine release dataset with MHC class I information, we utilized the PRIME-1.0 and PRIME-2.0 datasets (*22*). The PRIME datasets were preprocessed following the same methodology as described by (*11*). Then the dataset underwent prefiltering and removal of overlapping sequences with the benchmark set as outlined in Section A scrupulous data curation and leakage control strategy. We note that we did not allow any overlapping with the benchmark set in either the training or validation set, applying the stringent filtering method as described in Section Sequence identity filtering using CD-HIT. This process resulted in 1,512 positive and 5,163 negative data points, totaling 6,675. The dataset was split into training and validation sets in a 9:1 ratio, ensuring that peptides in the training set did not overlap with those in the validation set using the stringent filtering described in Section Sequence identity filtering using CD-HIT.

For the task IM-pep, a cytokine release dataset without MHC information, we utilized the datasets from (*21*). Since the number of negative samples was exceedingly high, we assumed that using all negatives would impair the model’s ability to learn a meaningful positive signal, so we randomly sampled 1,000 negative sequences. Then the dataset underwent prefiltering and removal of overlapping sequences with the benchmark set as outlined in Section A scrupulous data curation and leakage control strategy. We note that we did not allow any overlapping with the benchmark set in either the training or validation set, applying the stringent filtering method as detailed in Section Sequence identity filtering using CD-HIT. This process resulted in 142 positive and 996 negative samples, totaling 1,138. The dataset was split into training and validation sets in a 9:1 ratio, ensuring that peptides in the training set did not overlap with those in the validation set using the stringent filtering described in Section Sequence identity filtering using CD-HIT.

For the task IM-II, a cytokine release dataset with MHC class II information, we used datasets from IEDB (*12*) and MHCBN (*20*) to gather data on the immunogenicity of peptide-HLA class II pairs. These datasets were preprocessed following the methodology described by (*45*), resulting in 2,956 positive and 2,013 negative data points, totaling 4,969. The model was trained on this dataset using a binary classification task. The dataset was split into training and validation sets in a 9:1 ratio, ensuring that peptides in the training set did not overlap with those in the validation set using the stringent filtering method as outlined in Section Sequence identity filtering using CD-HIT.

### Cramer’s V analysis of label concordance

To evaluate the association between labels from datasets representing intermediate biological processes (e.g., peptide–MHC binding, TCR–pMHC interaction) and immunogenicity labels (cytokine release), we computed *Cramer’s V* statistics based on contingency tables constructed from overlap-ping peptides.

Peptide pairs with exact match of 9-mer fragment between datasets were identified, and the corresponding binary labels from the source (training) and target (test) datasets were extracted. For each dataset group defined by the training task identifier, we generated a 2 × 2 contingency table by cross-tabulating the number of peptides with each combination of train and test labels. To ensure uniformity across groups, the contingency table was reindexed to explicitly include all possible label pairs (0, 1), with missing entries filled as zeros.

Cramer’s V was calculated using the formula:

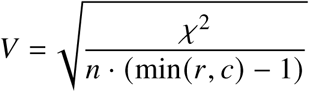

where χ^2^ is the chi-square statistic obtained from the contingency table, *n* is the total number of overlapping peptide pairs, and *r*, *c* are the number of rows and columns, respectively. The associated *p*-value was derived from the chi-square test using SciPy’s chi2 contingency function.

This analysis was performed both on the combined dataset and on subsets corresponding to individual training tasks, each representing distinct biological processes such as p-MHC binding affinity, p-MHC elution, or TCR interaction. The results were visualized using heatmaps to display the distribution of label concordance.

In the overall dataset, we observed a weak but statistically significant association (Cramer’s *V* = 0.0339, *p* = 0.0265), suggesting the presence of minor but informative signals between intermediate biological steps and immunogenicity. These results support the hypothesis that, despite label inversion, weak signals embedded in upstream datasets may contribute to the development of more robust immunogenicity prediction models.

### Sequence identity filtering using CD-HIT

To rigorously prevent data leakage between training and benchmark datasets, we applied a peptide-level identity filtering strategy using CD-HIT (*46*), based on 9-mer fragment identity. This process was designed under the assumption that the immunogenic signal of a peptide is often localized within a core 9-mer.

All peptides from both the training and benchmark sets were processed into overlapping 9-mer fragments using a sliding window approach. For peptides shorter than 9 residues, the full-length sequence was retained as a single fragment. Each fragment was assigned a unique label (e.g., train 0 0, test 1 3) and saved in FASTA format. The combined set of training and test 9-mers was clustered using CD-HIT with a sequence identity threshold of 0.8 (-c 0.8, to ensure less than 2 amino acids mismatches is tolerated for 9-mer fragment) and word size of 2 (-n 2).

From the resulting cluster assignments, we identified training peptides as overlapping if any of their 9-mer fragments were clustered together with a fragment derived from the benchmark set. These peptides were excluded from the final training set to eliminate potential memorization or indirect data contamination.

This strict filtering step ensured that the training set contained no sequences with high similarity to any benchmark peptide at the 9-mer level. The overall procedure is illustrated in Fig. 3B.Fig. 3B.

### Model Architecture

T-SCAPE employs a convolutional neural network (CNN)-based model to represent protein amino acid sequences, following the encoder and decoder architecture proposed by (*47*). While most protein language models utilize transformers (*48, 49*) or LSTMs (*11, 50*), we chose a CNN-based model. This decision aligns with the characteristics of T-cell immunogenicity tasks, where long protein sequences are fragmented into 9-mer peptides. This eliminates the need for global sequence attention and instead emphasizes local peptide features. We utilized kernel size 3 to catch the local feature of peptides. Similar approaches have been successfully adopted in various methods (*51,52*), demonstrating that CNN-based models can perform comparably well to transformer or LSTM-based architectures.

We employed the ByteNet architecture, a CNN-based encoder-decoder framework, as the core components of our model. We utilized the ByteNet encoder architecture to model the integrative encoder and task-specific encoder, except for conditioned sequence masked region prediction tasks. For the tasks Hu-MLM and EL, we used the conditioned ByteNet Decoder, which was slightly adjusted by adding an attention module to incorporate the condition into the embedding. For other tasks, we used the conventional ByteNet Decoder. As depicted in Figure 2, the embeddings generated by ByteNet were utilized as queries and values in the attention network, with each condition acting as the key.

By combining these components, we designed the integrative encoder, task-specific encoder/decoder, as shown in Figure 2. The integrative encoder consists of three ByteNet models that embed peptide, MHC class I pseudo sequence, and MHC class II pseudo sequence. These embeddings are combined using a gated attention mechanism which calculates attention scores as described in Equation 3. These scores are then used as weights to compute a weighted sum of the embeddings, as shown in Equation 2. For tasks that do not require MHC pseudo sequence (e.g., Hu-CLS), the output of the integrative encoder defaults to the peptide embedding.

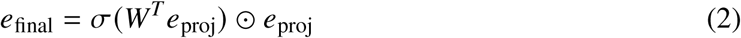

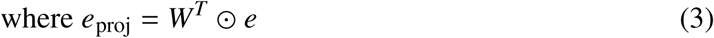

This gated attention mechanism is also applied to the task-specific encoder when multiple inputs are needed. The embeddings from the integrative encoder and the task-specific encoder are then concatenated and fed into the task-specific decoder, which is a simple MLP layer.

Lastly, the framework includes a discriminator for adversarial training of integrative encoder. The discriminator is implemented as a simple MLP layer that takes the integrative encoder embedding as input and outputs the data source classification probability. This component enables adversarial learning, where the model is trained using a gradient reversal mechanism to improve robustness and generalization.

### Training scheme

We applied a multi-domain adversarial learning scheme (*53*), a method widely used in other biomedical applications (*54, 35*). The training of T-SCAPE involved three types of loss functions: task-specific loss, orthogonal loss, and adversarial loss.

#### Task-specific Loss

For each task, we used task-specific loss as described in Equations (4) and (5).

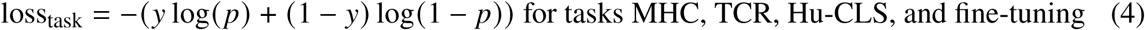

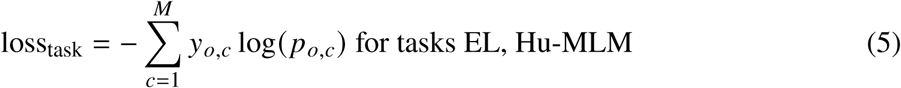

For binary classification tasks (e.g., MHC, TCR, and Hu-CLS), we used binary cross-entropy loss as in Equation (4), where *y* represents the label and *p* the predicted probability. For sequence masked region prediction tasks (e.g., EL and Hu-MLM), we employed cross-entropy loss to predict masked and mutated amino acids, as shown in Equation (5). In this context, *c* represents tokens, including amino acid codes, as well as stop, gap, mask, and start tokens. Here, *p_o,c_* is the model’s probability for each token, and *y*_*o*,*c*_ denotes the correct token. We masked 80% and mutated 10% of the sequence, following (*37*).

#### Orthogonal Loss

To differentiate the task-specific encoder output from the integrative encoder, we applied an orthogonal loss to make the embeddings orthogonal, as described in Equation (6).

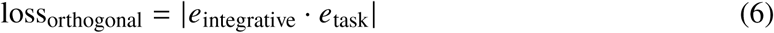

In this equation, *e*_integrative_ represents the output embedding from the integrative encoder, and *e*_task_ denotes the output embedding from the task-specific encoder.

#### Adversarial loss

To ensure the integrative encoder captures common information across data sources, we applied cross-entropy loss to the discriminator, similar to Equation (5), and adversarial loss to the integrative encoder, as defined in Equation (7).

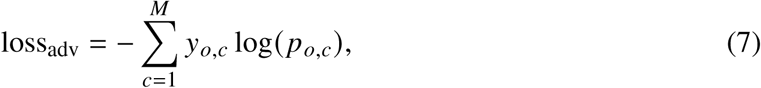

In this equation, *c* represents the task index, *p*_*o*,*c*_ is the model’s probability for each class, and *y*_*o*,*c*_ is the true class label.

The total loss was optimized by linearly combining these loss components, as described in Equation (8).

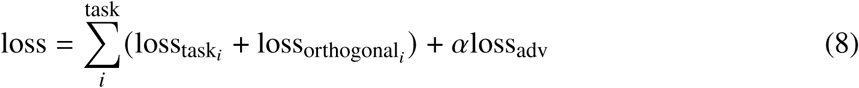

The coefficient α was set to 5 after testing values of 1, and 5. All models and training processes were implemented using PyTorch (*55*). The training utilized the Adam optimizer (*56*) and a cosine annealing learning rate scheduler (*57*) were employed, with a learning rate of 1e-5, a batch size of 40, gradient accumulation of 4, a model dimension of 280, and 6 layers, with a total training duration of 10 days on an RTX A6000 GPU.

For fine-tuning, we employed binary cross-entropy loss as described in Equaion 4, using the same optimizer and scheduler as in pretraining. The fine-tuning settings included a learning rate was of 5e-5, a batch size of 128, and a total of 100 epochs.

### Bootstrapping and statistical evaluation

To assess the reliability and statistical significance of each model’s performance, we applied a resampling-based evaluation framework using bootstrapping and non-parametric hypothesis testing.

#### Bootstrap confidence intervals

For each model and each metric—AUROC, PR-AUC, precision (PPV), recall, and mean PPVn—we generated 1,000 bootstrap replicates by sampling with replacement from the full test dataset to compute 95% confidence intervals. For each bootstrap sample, we computed the metric of interest, and used the resulting empirical distribution to estimate the 95% confidence interval via percentile estimation (2.5th to 97.5th percentiles). Metrics involving ranking (e.g., PPVn) were computed by sorting predictions and averaging top-*n* precision scores across the range of *n*.

#### Paired statistical testing

To assess whether performance differences between models were statistically significant, we used the Wilcoxon signed-rank test on the paired bootstrap distributions of each metric. Specifically, we compared T-SCAPE-IM to the models with the next lower and next higher performance ranks, as well as to its baseline variant, t-scape-im, to assess the significance of pairwise differences. To account for multiple comparisons across baseline models, we applied the Bonferroni correction to the resulting *p*-values.

#### Ablation analysis

The same bootstrapping and statistical testing procedure was applied to the ablation study. For each ablated variant of T-SCAPE-IM, we computed confidence intervals and conducted paired comparisons against the original model to quantify the contribution of each pretraining dataset and module. Performance differences and statistical significance were visualized to highlight the impact of each ablation.

#### Stratified bootstrap analysis

To evaluate model robustness across biologically meaningful subgroups, we additionally performed stratified bootstrapping based on HLA allele type (e.g., A, B, C for class I; DP, DQ, DR for class II) and peptide length (e.g., 8–11-mers). For each subgroup, bootstrap resampling was repeated within the subgroup, and the metric distributions were independently computed. Stratified results were visualized as line plots with shaded confidence intervals.

All statistical analyses and visualizations were implemented using Python, with the scikit-learn, scipy, and matplotlib libraries.

### Benchmark label

Since we defined all the tasks in T-SCAPE as classification tasks, continuous measurements and scoring outputs were preprocessed into binary classification labels or standardized probability-like values depending on the context of each dataset.

For the ADA (anti-drug antibody) evaluation dataset, immunogenicity values indicated how many individuals exhibited an ADA response to a given antibody. We assigned binary labels such that any non-zero immunogenicity value was labeled as 1 (immunogenic), while 0.0 was labeled as 0 (non-immunogenic). To enable fair comparison with other models, which predominantly output humanness or human-likeness scores, we transformed those scores to align with the immunogenicity scale.

Scores where higher values indicated human-likeness (e.g., AbNatiV, Hu-mAb, Germline content) were inverted by:

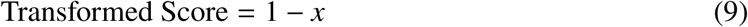

Percentile-based scores such as T20 were converted using:

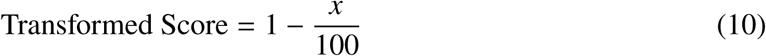

Z-score-based outputs (e.g., Z-score, IgReconstruct, AbLSTM) were standardized by applying the cumulative distribution function (CDF) of the standard normal distribution:

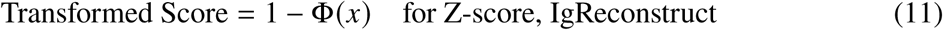

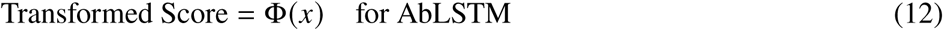

depending on the original directionality of the score, where Φ(*x*) denotes the standard normal CDF. Likewise, OASis identity scores (loose to strict), were transformed as:

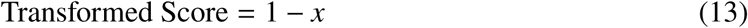

For benchmarking T-SCAPE-MHC, peptide-MHC binding affinity values (IC50) were converted into binary labels and normalized scores. Specifically, for entries where the “Measurement type” column indicated a non-binary value, the IC50 value was converted to a binary label: peptides with IC50 ≤ 500 nM were labeled as 1 (binder), and those with IC50 > 500 nM were labeled as 0 (non-binder). This threshold is commonly used to define peptide-MHC binding and IEDB guideline (*12*). In addition, for each MHC binding prediction model, predicted IC50 values were transformed using the following function:

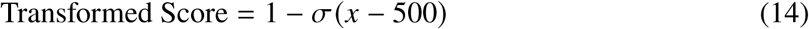

where σ is the sigmoid function and *x* is the raw IC50 value. This transformation ensures that stronger predicted binding is mapped to a score closer to 1, facilitating consistent comparison with binary classifiers. Non-numeric entries (e.g., “-”) were excluded from the transformation. The entire procedure was implemented in a Python script provided in the supplementary materials.

These label and score transformation procedures allowed all datasets and models to be consistently evaluated under a unified classification framework.

### Embedding quality benchmark

To evaluate the quality of the embeddings generated by the integrative encoder, we first converted them into 2D tensors using t-SNE (*58*), implemented in scikit-learn (*59*). The quality of these embeddings was then quantified using two widely recognized metrics: the Silhouette Score (SI) and Adjusted Mutual Information (AMI), which are commonly used to assess embedding and clustering quality (*60, 61, 62*). The SI is calculated using Equation (15).

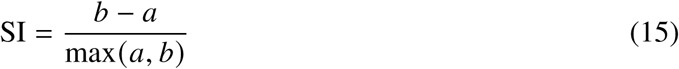

where a represents the mean intra-cluster distance, and b denotes the mean nearest-cluster distance. The AMI quantifies the mutual information by Equation (16).

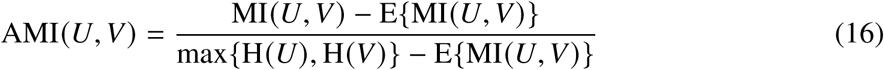

where MI(U,V) denotes mutual information between two distributions U and V, H(U) represents entropy of distribution U, and E(MI(U,V)) denotes expected mutual information between two distributions U and V.

## Author contributions

Jeonghyeon Kim, Jinsung Noh, and Chaok Seok conceived the study and conducted the experiments. Jeonghyeon Kim, Nuri Jung, Jayyoon Lee, Namhyuk Cho, Jinsung Noh, and Chaok Seok analyzed the results. Jeonghyeon Kim drafted the manuscript, while Nuri Jung implemented the web server. All authors—Jeonghyeon Kim, Nuri Jung, Jayyoon Lee, Namhyuk Cho, Jinsung Noh, and Chaok Seok—contributed to revising the manuscript.

## Data and materials availability

All datasets used for training, validation, and benchmarking of T-SCAPE are available at https://figshare.com/articles/dataset/TITANiAN_ benchmark_result/26323582. Data were sourced from publicly available databases, including IEDB (*12*), VDJdb (*24*), MCPAS-TCR (*39*), ImmuneCODE (*40*), TBAdb (*41*), 10X Genomics (*42*), OAS (*63*), UNIPROT (*16*), PRIME (*23*), MHCBN (*20*), BigMHC (*11*), Biophi (*19*),and Panpep (*44*).

The T-SCAPE code repository, which includes the model architecture, training code, inference code, and analysis code, is available at https://github.com/seoklab/T-SCAPE. Additionally, a user-friendly web server is provided for easy access and utilization of the model at https://galaxy.seoklab.org/design/t-scape.

## Supplementary materials

Supplementary Text

Figs. S1 to S8

Tables S1 to S4

References (*12-63)*

