## Supplemental_file for "T-SCAPE: T-cell Immunogenicity Scoring via Cross-domain Aided Predictive Engine"

**This PDF file includes:**

Supplementary Text

Figures S1 to S8

Tables S1 to S4

### Supplementary Text

#### Dataset curation and composition

A rigorous data curation and partitioning process was implemented to ensure a fair and robust evaluation of the model's generalization capabilities by preventing data leakage. Figure S1 provides a comprehensive summary of this process.

Figure S1A and B detail the statistics for our pretraining and fine-tuning datasets, respectively. They list the initial number of data points and the final counts after applying our benchmark-based clustering filter, which removes sequences with high homology to any benchmark set. The data was subsequently partitioned into leakage-free training (90%) and validation (10%) sets.

The heatmap in Figure S1C provides a comprehensive visualization of the 9-mer overlap analysis that guided our data curation process. The values shown represent the number of shared 9-mer sequences between any two datasets before the final filtering. The numbers highlighted in bold are of particular importance, as they show the initial overlaps identified between our training/validation data and the independent benchmark sets. To prevent data leakage and ensure a fair evaluation, all of these bolded, overlapping sequences were subsequently removed from their respective training and validation sets. This critical filtering step guarantees that the model's performance on the benchmark tasks is a true measure of its generalization capabilities.

#### Method representation of each models

We demonstrate the application and performance of our T-SCAPE framework on several key immunological prediction tasks, as illustrated in Figure S2. The model's modularity permits task-specific configurations. Specifically, for T-cell epitope immunogenicity and ADA level prediction, the T-SCAPE-IM module employs the output from the integrative encoder with the Task-IM decoder (Figure S2A-B). For peptide-MHC and TCR-peptide-MHC binding predictions, the T-SCAPE-MHC and T-SCAPE-TCR modules, respectively, concatenate features from the shared integrative encoder with those from a task-specific encoder prior to processing by the final decoder (Figure S2C-D). A comprehensive performance evaluation of these configurations is presented across six distinct benchmarks, including neoantigen discovery, infectious disease vaccines, FDA-approved ADA levels, and IEDB MHC class I/II binding (Figure S2E-J). The results underscore the model's robust performance and versatility across these diverse predictive challenges.

#### Overlap analysis for antibody case study sequences

The extent of sequence overlap for the de-immunized versions with our training tasks is detailed in Supplementary Table S1. This analysis confirms that the 9-mer peptides from the case study sequences have minimal overlap with our various training datasets, reinforcing the validity of the evaluation. Additionally, the number of samples (N) in the score distribution plots of Figure 7 corresponds to the total number of unique 9-mer peptides for each sequence, as listed in the table.

#### T-SCAPE-MHC: Predicting peptide-MHC binding

Predicting peptide-MHC interactions is a crucial initial step in identifying potential immunogenic peptides for applications such as cancer immunotherapy and vaccine development. While the T-cell epitope immunogenicity prediction results presented in Figures4 A and B demonstrate the utility of *in silico* methods, they also highlight that these approaches are not yet infallible, underscoring the need for complementary strategies. One such strategy involves pre-filtering peptide candidates based on predicted MHC binding affinity using models trained on peptide-MHC interaction data. This pre-selection process can significantly reduce the search space for downstream immunogenicity assays. Therefore, in the context of peptide-MHC binding prediction, Precision and Mean PPVn are considered key metrics for evaluating the accuracy of identifying true binders. To ensure that high Precision and Mean PPVn are not achieved at the expense of Recall, we also evaluate the PR-AUC. Additionally, AUROC is included as a general measure of the classifier's overall performance in distinguishing between binders and non-binders.

As depicted in Figure S2C, T-SCAPE-MHC utilizes the shared integrative encoder alongside the dedicated Task-MHC encoder and decoder. T-SCAPE-MHC operates solely on the parameters learned during the pretraining phase, without undergoing any additional task-specific fine-tuning.

To evaluate the performance of T-SCAPE-MHC I and II in predicting peptide-MHC binding, we employed the IEDB benchmark set (12) with data collected after December 25, 2020 (for MHC class I: size = 624, positives = 344, negatives = 280) and after February 19, 2021 (for MHC class II: size = 723, positives = 242, negatives = 481). Consistent with our rigorous data leakage prevention strategy, we excluded any training sequence that shared 9-mer fragments with less than

2 mismatches with any peptide in these benchmark sets. This stringent filtering ensures that the reported performance reflects genuine generalization capabilities rather than the memorization of highly similar sequences. Furthermore, we carefully curated the benchmark sets to encompass a diverse range of HLA alleles. Notably, the class I benchmark includes the HLA-E allele (HLA A - 368, B - 84, E - 172) to specifically assess each model's ability to generalize to alleles that are underrepresented in the training data.

To rigorously evaluate the statistical significance of T-SCAPE-MHC's performance gains, we employed bootstrap analysis with 1,000 resamples to estimate 95% confidence intervals for each evaluation metric. Additionally, we conducted paired Wilcoxon signed-rank tests with Bonferroni correction to assess the statistical significance of the observed differences. Specifically, we compared T-SCAPE-MHC against the models ranked immediately below and above it in performance to pinpoint the significance of improvements relative to its closest competitors.

As explicitly noted in Figure S3, the T-SCAPE-MHC models were not fine-tuned for this task, and their model architecture and parameters differ from those of T-SCAPE-IM. This evaluation, therefore, assesses the performance derived solely from knowledge acquired during pretraining.

The performance of T-SCAPE-MHC I in predicting MHC class I binding was benchmarked against a panel of established methods. As illustrated in Figure S3A, it achieves a Mean PPVn of 0.770, which was marginally lower, a statistically significant difference, compared to IEDB Consensus (0.804), but showed improvements over models with lower performance ($p < 0.001$). Although its precision (0.619) was significantly lower than that of NetMHCpan 4.0 EL (0.775), its PR-AUC (0.725) was significantly higher than that of NetMHCpan 4.0 EL, indicating a favorable balance between precision and recall ($p < 0.001$). The AUROC of T-SCAPE-MHC I (0.674) was marginally lower, a statistically significant difference, compared to NetMHCpan 4.1 EL (0.691) but surpassed several earlier models ($p < 0.001$).

Figure S3B presents a performance comparison of T-SCAPE-MHC II against several established MHC class II binding prediction methods. It achieves a Mean PPVn of 0.613, which was marginally higher, a statistically significant difference, compared to NN-align 2.3 (0.598; $p < 0.001$) and ranked third among the compared methods. While its precision (0.349) and PR-AUC (0.521) were significantly lower than those of the leading models such as NetMHCIIpan-4.0 EL (precision: 1.000), T-SCAPE-MHC II achieved a marginally higher precision, which was statistically significant, compared to IEDB Consensus (0.338; $p < 0.001$). Its AUROC (0.657) was significantly lower than that of NetMHCIIpan-4.0 EL (0.817) but significantly higher than Tepitope (0.500; $p < 0.001$). The relatively larger performance gap observed in MHC class II prediction may be attributed to the inherent variability in peptide lengths and binding modes associated with MHC class II molecules.

#### T-SCAPE-TCR: Predicting TCR-peptide-MHC binding

Predicting peptide-MHC-TCR binding holds significant potential for applications such as rational TCR design in Adoptive Cell Transfer (ACT)-based therapy and the analysis of infectious disease cohorts. (44) For these applications, the ability to accurately identify high-confidence interactions is crucial, making Precision and Mean PPVn key performance metrics. To provide a comprehensive evaluation of the classifier's capabilities, we also assess PR-AUC and AUROC. In contrast to the immunogenicity prediction task discussed in Section T-SCAPE-IM: T-cell epitope immunogenicity prediction for specific peptide-MHC pairs, the relatively balanced distribution of positive and negative samples in this dataset allows us to emphasize AUROC as a similarly important metric alongside PR-AUC.

T-SCAPE-TCR, which comprises the shared integrative encoder and the dedicated Task-TCR encoder/decoder (as illustrated in Figure S2D), predicts TCR-peptide-MHC binding by leveraging solely the parameters learned during the pretraining phase, without any task-specific fine-tuning. We evaluated T-SCAPE-TCR on two distinct benchmark sets: the Panpep Zero-shot test set (size: 786, positive: 393, negative: 393) and the ACT test set (size: 26, positive: 13, negative: 13) (44). While we acknowledge the inherent limitations of the ACT test set due to its small size, its data provides a uniquely valuable benchmark for clinical applications. This high relevance stems from its focus on tumor-specific neoantigens—the precise targets of ACT—and its use of TCRs derived from patients with metastatic cancer, thus reflecting a real-world therapeutic scenario. To assess its performance, T-SCAPE-TCR was compared against several task-specific methods, including Pan-pep (44), DLpTCR (14), pMTnet (64), and ERGO2.

As shown in Figure S3C, our evaluation on the zero-shot test set revealed that T-SCAPE-TCR's performance was significantly lower than Pan-pep's but significantly outperformed all other methods. It attained a Mean PPVn of 0.656, which was significantly lower than Pan-pep (0.801) but significantly higher than the third-best model, ERGO2 (0.500; $p < 0.001$). A similar trend was observed in Precision, where its score of 0.650 was significantly lower than Pan-pep's (0.717) but significantly exceeded DLpTCR's (0.505; $p < 0.001$). Its PR-AUC of 0.709 was marginally lower, a statistically significant difference, compared to Pan-pep (0.747), while significantly surpassing ERGO2 (0.500; $p < 0.001$). The AUROC of T-SCAPE-TCR (0.630) was also significantly lower than Pan-pep's (0.739) but exceeded all other models, with significant differences observed (vs. DLpTCR: 0.504; $p < 0.001$).

On the ACT-based therapy discovery set (Figure S3D), T-SCAPE-TCR demonstrated a highly competitive performance. It achieved a Mean PPVn of 0.800, which was significantly lower than that of Pan-pep (0.883) but significantly higher than DLpTCR (0.621; $p < 0.001$). Its Precision was identical to Pan-pep (0.778) and significantly higher than all other methods ($p < 0.001$). The model's PR-AUC (0.745) was significantly lower than Pan-pep's (0.819) but significantly higher than DLpTCR's (0.595; $p < 0.001$). Notably, T-SCAPE-TCR achieved the highest AUROC among all evaluated methods on this dataset, with its score of 0.769 representing a marginally higher performance, which was statistically significant, compared to Pan-pep's 0.751 ($p < 0.001$). These competitive results are noteworthy given that Task-TCR is a lightweight, auxiliary module. The model prioritizes the pMHC representation and incorporates minimal TCR-specific processing, yet still achieves reasonable performance on TCR-centric tasks.

Furthermore, we recognized potential concerns regarding negative sampling bias in TCR specificity prediction. A recent study by (65) suggested that randomly shuffling the positive set to generate negative samples is a more appropriate, less biased method for Pan-pep's test set. Acknowledging this, we applied a similar approach to our TCR zeroshot evaluation. This re-evaluation resulted in a performance decrease for our model to an AUROC of 0.511, PR-AUC of 0.624, Precision of 0.508, and Mean PPVn of 0.546. However, our revised AUROC of 0.511 is still slightly higher than the previously reported top performance of Pan-pep (0.497), suggesting that T-SCAPE-TCR is less susceptible to this particular form of evaluation bias.

#### ROC curve for benchmark results

Figure S4 (left) displays the Receiver Operating Characteristic (ROC) curve for the benchmark detailed in the main text, with the navy line indicating the ROC curve of a random classifier as a reference for each model's performance. Figure S5 (right) presents the Precision-Recall curve for the benchmarks mentioned in the main text. From the top, the curves represent: T-SCAPE-IM benchmark result for Neoantigen discovery set," T-SCAPE-IM benchmark result for Infectious disease set," T-SCAPE-IM benchmark result for ADA level test," T-SCAPE-MHC I benchmark result for IEDB benchmark set," T-SCAPE-MHC II benchmark result for IEDB benchmark set," T-SCAPE-TCR benchmark result for zero-shot set," and "T-SCAPE-TCR benchmark result for ACT-based therapy discovery set."

#### PPVn based on the number of top predictions

Figure S6 depicts the line plot of PPV vs. Top N, with a dashed parallel line representing the random classifier to provide a reference for each model's PPV vs. Top N curve. From the top left, the curves represent: T-SCAPE-IM benchmark result for Neoantigen discovery set," T-SCAPE-IM benchmark result for Infectious disease set," T-SCAPE-MHC I benchmark result for IEDB benchmark set," T-SCAPE-MHC II benchmark result for IEDB benchmark set," T-SCAPE-TCR benchmark result for zero-shot set," and T-SCAPE-TCR benchmark result for ACT-based therapy discovery set."

#### Optimizing the coefficient for the adversarial loss and orthogonality constraint for encoder specialization

To determine the adversarial loss coefficient $\alpha$, we conducted experiments to evaluate how adjusting $\alpha$ affects the performance of our final model, T-SCAPE-IM. We pretrained each model under identical conditions, varying only $\alpha$, and monitored the validation loss during the fine-tuning steps. As depicted in Figure S7, a larger $\alpha$ value results in lower validation loss and higher validation set AUROC, indicating that a larger $\alpha$ value enhances the model's generalizability in predicting immunogenicity. We attribute this to the fact that a higher adversarial loss coefficient enforces more rigorous alignment from different data sources in the integrative encoder.

To ensure that our model effectively disentangles shared and task-specific features, we introduced an orthogonality constraint between the shared encoder and the various task-specific encoders. The motivation stems from a conceptual Venn diagram of multi-task information: a central region of shared features useful for all tasks, and unique, non-overlapping regions of features specific to each individual task. By penalizing the similarity (via cosine similarity) between the outputs of the shared and task-specific encoders, we encourage them to specialize. The effectiveness of this approach is demonstrated in Figure S7B, where the orthogonal loss consistently decreases during training, confirming that the learned representations become more distinct and specialized.

### t-SNE plot of T-SCAPE-IM embedding on various datasets

We also generated a t-SNE plot of the test set embeddings on vairous test set, shown in Figure S8, to demonstrate that T-SCAPE-IM's performance is not solely dependent on the decoder's classification but is also influenced by the effective embeddings from pretraining and fine-tuning. The embeddings were taken from the output tensor of the integrative encoder on neoantigen discovery set and infectious disease vaccine discovery set. According to figure S8A), for the neoantigen discovery set, the highlighted region of the figure shows that immunogenic peptides cluster with minimal non-immunogenic peptide points. Additionally, the Silhouette score (SI) and Adjusted Mutual Information (AMI) values are 0.132 and 0.017, respectively, indicating that the embedding is well-clustered compared to random embeddings, which have SI values of -1.0 and AMI values of 0.0. These analyses suggest that T-SCAPE-IM's embeddings are effective in discriminating between immunogenic and non-immunogenic peptides. Moreover, the highlighted region of the figure S8B) , which is t-SNE plot on infectious disease vaccine discovery set, shows that non-immunogenic peptides cluster with minimal immunogenic peptide points. The Silhouette score (SI) and Adjusted Mutual Information (AMI) values are 0.203 and 0.045, respectively, indicating that the embedding is well-clustered compared to random embeddings, which have SI values of -1.0 and AMI values of 0.0. These analyses suggest that T-SCAPE-IM's embeddings are effective in discriminating between immunogenic and non-immunogenic peptides also in infectious disease vaccine discovery set.

Figure S8C) shows the t-SNE plot of T-SCAPE-IM's embeddings for the ADA benchmark set. To obtain the embedding of each antibody drug sequence, we applied max pooling to each peptide's sequence. Since immunogenicity can be triggered by a single problematic sequence, max pooling was used to derive the protein's overall embedding. The highlighted region of the figure demonstrates that T-SCAPE-IM effectively creates meaningful embeddings regarding the immunogenicity of protein sequences. The Silhouette score (SI) and Adjusted Mutual Information (AMI) values are 0.057 and 0.045, respectively, indicating that the embedding is well-clustered compared to random embeddings, which have SI values of -1.0 and AMI values of 0.0.This indicates that T-SCAPE's performance does not solely depend on the decoding classifier.

### Overview of the Dataset Composition

**A** Pretraining Set

| Task | Initial set | After benchmark clustering filter | Leakage-free training set (90%) | Leakage-free validation set (10%) |
|---|---|---|---|---|
| Task-Hu | 8,629,139 | 7,965,334 | 7,169,643 | 795,691 |
| Task-MHC BA I | 130,065 | 129,967 | 117,435 | 12,532 |
| Task-MHC EL I | 557,161 | 557,151 | 501,640 | 55,511 |
| Task-MHC BA II | 516,482 | 516,460 | 464,966 | 51,494 |
| Task-MHC EL II | 206,059 | 206,055 | 185,602 | 20,453 |
| Task-TCR | 130,111 | 130,060 | 128,720 | 1,340 |
| Total | 10,169,017 | 9,505,027 | 15,736,256 | 1,734,105 |

**B** Fine-tuning Set

| Task | Initial set | After benchmark clustering filter | Leakage-free training set (90%) | Leakage-free validation set (10%) |
|---|---|---|---|---|
| Task-IM (w/ MHC I) | 6,681 | 6,675 | 6,029 | 646 |
| Task-IM (w/ MHC II) | 7,314 | 7,077 | 6,294 | 783 |
| Task-IM (w/o MHC) | 1,138 | 1,138 | 1,019 | 119 |
| Total | 15,034 | 14,890 | 13,342 | 1,548 |

### 9-mer Overlap between Datasets

**C**

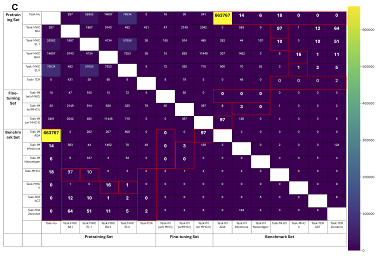

**Figure S1**: **Overview of dataset composition and 9-mer overlap analysis.** (**A**) Detailed counts for the pretraining datasets, showing the initial number of samples, the count after applying the benchmark clustering filter, and the final split into leakage-free training (90%) and validation (10%) sets. (**B**) Detailed counts for the fine-tuning datasets, following the same structure as (**A**). (**C**) A heatmap illustrating the 9-mer peptide sequence overlap between all datasets. Off-diagonal elements quantify the number of shared 9-mers between any two sets. The numbers shown in bold and red boxes represent the initial overlap between the training/validation sets and the benchmark sets; these specific sequences were removed during our curation process to prevent data leakage. The diagonal is left blank as the analysis focuses on inter-dataset comparisons.

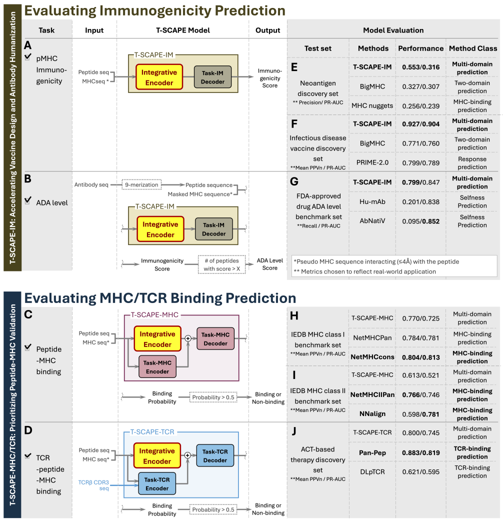


**Figure S2**: **T-SCAPE module usage for prediction tasks (A to D) and performance comparison across benchmarks (E to J).** T-SCAPE utilizes different module configurations for specific tasks. (**A**) For T-cell epitope immunogenicity prediction and (**B**) ADA level prediction, T-SCAPE-IM uses the integrative encoder output with the Task-IM decoder. (**C**) For peptide-MHC binding (T-SCAPE-MHC) and (**D**) TCR-peptide-MHC binding (T-SCAPE-TCR), features from the shared integrative encoder and the respective task-specific encoder are concatenated before being processed by the task-specific decoder. Performance summaries are presented for T-SCAPE-IM assessed on (**E**) the neoantigen discovery set, (**F**) the infectious disease vaccine discovery set, and (**G**) the FDA-approved ADA level benchmark; T-SCAPE-MHC evaluated on (**H**) the IEDB MHC class I and (**I**) IEDB MHC class II benchmarks; and T-SCAPE-TCR tested on (**J**) the ACT-based therapy discovery set. Performance metrics appropriate for each benchmark task are specified within the panels.

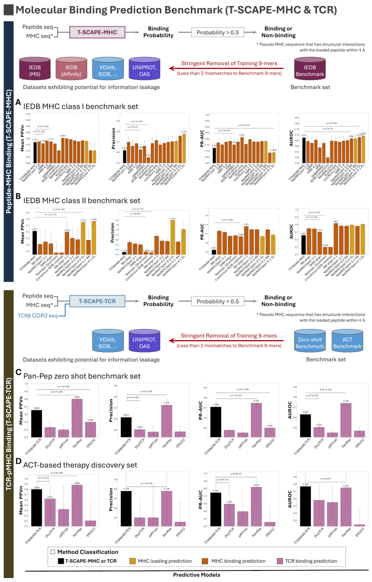


**Figure S3**: **Performance of pre-trained T-SCAPE auxiliary modules on binding prediction benchmarks.** Comparison of T-SCAPE-MHC I (**A**), T-SCAPE-MHC II (**B**), and T-SCAPE-TCR (**C**, **D**)—utilized solely with parameters learned during multi-domain pretraining without task-specific fine-tuning—against established task-specific binding prediction methods. Evaluations were performed on benchmarks specific to each task: (**A**) IEDB MHC class I binding (using data post 2020-12-25), (**B**) IEDB MHC class II binding (post 2021-02-19), (**C**) PanPep Zero-shot TCR binding, and (**D**) ACT-based therapy discovery TCR binding. Performance was assessed using Mean PPVn, Precision, PR-AUC, and AUROC. All metrics are presented with 95% confidence intervals (bootstrapping); statistical significance (p-values) was evaluated using paired Wilcoxon signed-rank tests with Bonferroni correction.

- **T-SCAPE-IM Benchmark result for Neoantigen discovery set**

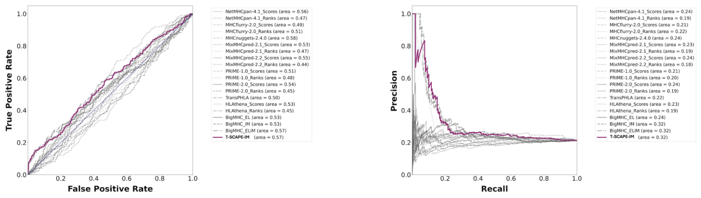


- **T-SCAPE-IM Benchmark result for Infectious disease set**

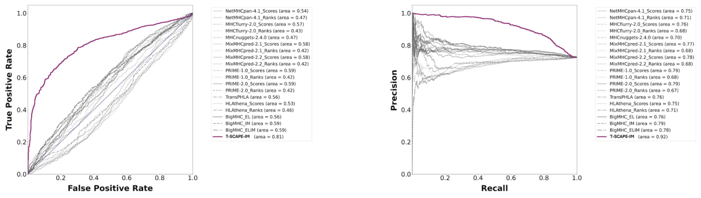


- **T-SCAPE-IM Benchmark result for ADA level set**

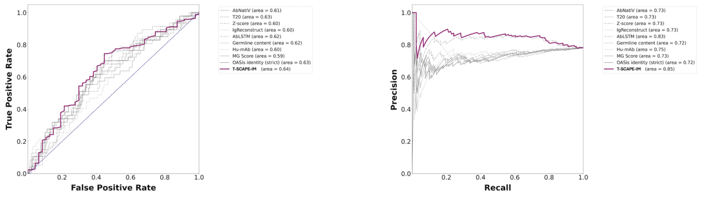


**Figure S4**: Left) Receiver Operating Characteristic (ROC) curve for the benchmarks described in the main text. The navy line represents the ROC curve of a random classifier, providing a reference for each model's ROC curve performance. Right) Precision-Recall curve for the benchmarks described in the main text. From the top, each curve represents: "T-SCAPE-IM benchmark result for Neoantigen discovery set", "T-SCAPE-IM benchmark result for Infectious disease set", and "T-SCAPE-IM benchmark result for ADA level test".

- **T-SCAPE-MHC I Benchmark result for IEDB benchmark set**

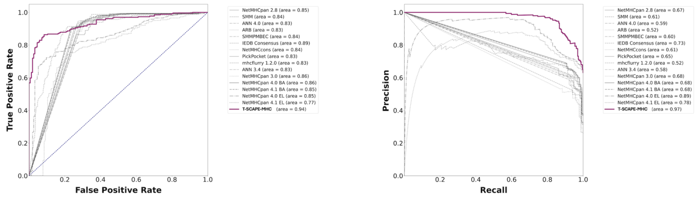


- **T-SCAPE-MHC II Benchmark result for IEDB benchmark set**

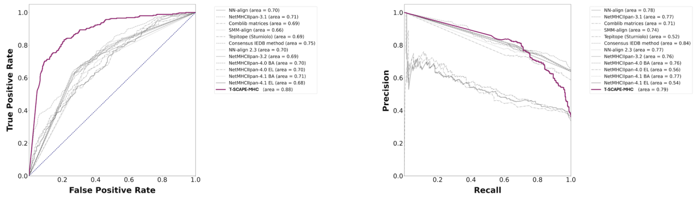


- **T-SCAPE-TCR Benchmark result for Zero shot set**

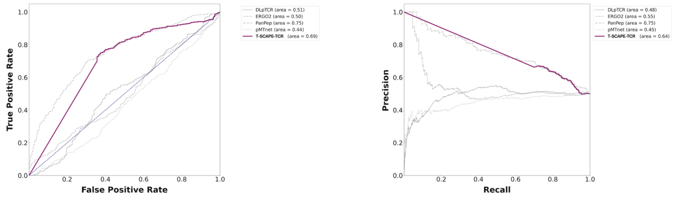


- **T-SCAPE-TCR Benchmark result for ACT-based therapy discovery set**

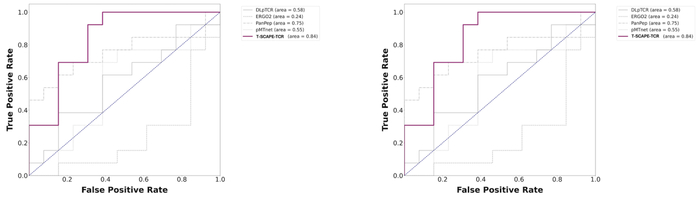


**Figure S5**: Left) Receiver Operating Characteristic (ROC) curve for the benchmarks described in the main text. The navy line represents the ROC curve of a random classifier, providing a reference for each model's ROC curve performance. Right) Precision-Recall curve for the benchmarks described in the main text. From the top, each curve represents: "T-SCAPE-MHC I benchmark result for IEDB benchmark set", "T-SCAPE-MHC II benchmark result for IEDB benchmark set", "T-SCAPE-TCR benchmark result for zero-shot set", and "T-SCAPE-TCR benchmark result for ACT-based therapy discovery set"

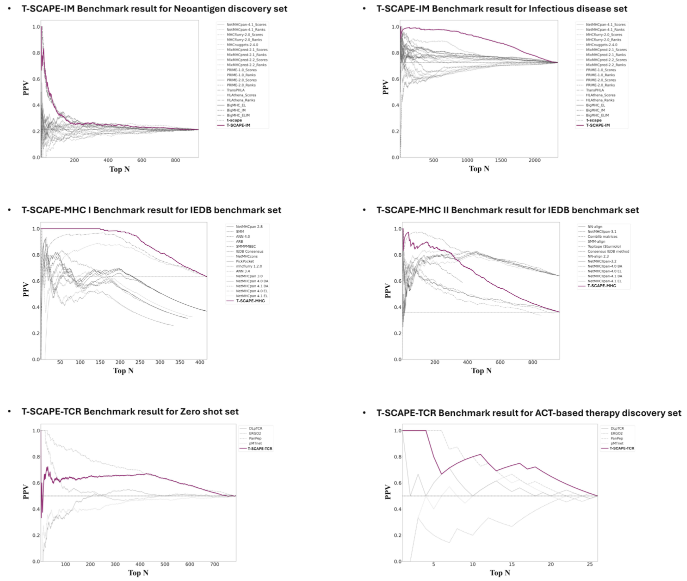


**Figure S6**: Line plot of PPV vs Top N. A dashed parallel line represents the random classifier, providing a reference for each model's PPV vs. Top N curve. From the top left, each curve represents: "T-SCAPE-IM benchmark result for Neoantigen discovery set", "T-SCAPE-IM benchmark result for Infectious disease set", "T-SCAPE-MHC I benchmark result for IEDB benchmark set", "T-SCAPE-MHC II benchmark result for IEDB benchmark set", "T-SCAPE-TCR benchmark result for zero-shot set", and "T-SCAPE-TCR benchmark result for ACT-based therapy discovery set".

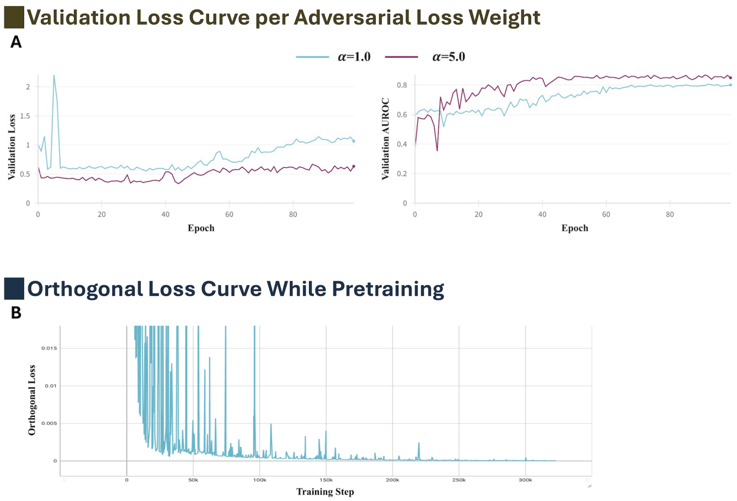


**Figure S7**: (**A**) Left) Epoch vs. Validation Loss curve, and Right) Epoch vs. Validation set AUROC curve in fine-tuning steps, depicting the impact of varying the adversarial loss coefficient $\alpha$ in the pretraining step. In both figures, higher $\alpha$ values result in improved performance, leading to the selection of $\alpha = 5.0$. (**B**) The plot displays the orthogonal loss throughout the training process, showing both the fluctuating raw loss and a smoothed curve overlaid to highlight the general trend. The consistent decrease indicates that the representations from the shared encoder and the task-specific encoders are becoming increasingly orthogonal. This enforces the specialization of the encoders to learn either common or task-specific features.

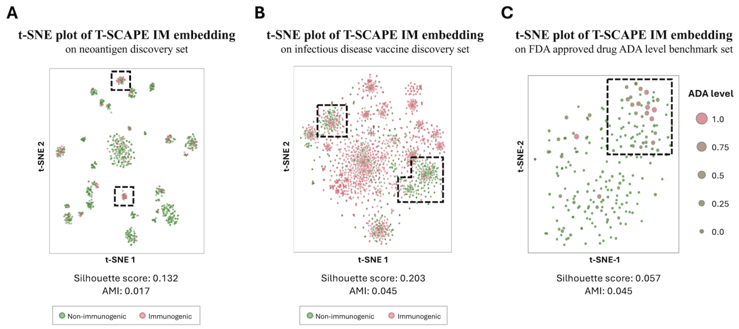


**Figure S8**: t-SNE plot of TIANiAN-IM embedding on various datasets. (**A**) A t-SNE plot of our embeddings for the neoantigen discovery set illustrates T-SCAPE-IM's effectiveness in distinguishing between immunogenic and non-immunogenic peptides. (**B**) A t-SNE plot of our embeddings for the infectious disease benchmark set illustrates T-SCAPE-IM's effectiveness in distinguishing between immunogenic and non-immunogenic peptides. (**C**) A t-SNE plot of our embeddings for the ADA level benchmark set illustrates T-SCAPE-IM's effectiveness in distinguishing between high and low ADA level antibody drug sequences.

**Table S1**: **Overlap analysis of unique 9-mer peptides from de-immunized antibody sequences used in the case study (Figure 7 G to J).** The table quantifies the number of shared 9-mers between each antibody chain and the various pretraining and fine-tuning datasets.

| | **Bevacizumab** | | **Etaracizumab** | | **Omalizumab** | | **Pembrolizumab** | |
|---|---|---|---|---|---|---|---|---|
| | **VH** | **VL** | **VH** | **VL** | **VH** | **VL** | **VH** | **VL** |
| Number of Unique 9-mer Peptide | 115 | 99 | 109 | 99 | 113 | 103 | 112 | 103 |
| Task-Hu | 10 | 3 | 7 | 3 | 1 | 11 | 8 | 2 |
| Task-MHC BA I | 0 | 0 | 0 | 0 | 0 | 0 | 0 | 0 |
| Task-MHC EL I | 0 | 2 | 2 | 1 | 13 | 12 | 0 | 13 |
| Task-MHC BA II | 0 | 2 | 1 | 0 | 6 | 3 | 0 | 3 |
| Task-MHC EL II | 0 | 4 | 2 | 0 | 3 | 4 | 0 | 1 |
| Task-TCR | 0 | 0 | 0 | 0 | 0 | 0 | 0 | 0 |
| Task-IM MHC I | 0 | 1 | 0 | 0 | 2 | 1 | 0 | 0 |
| Task-IM MHC II | 0 | 2 | 0 | 0 | 0 | 0 | 0 | 1 |
| Task-IM w/o MHC | 0 | 1 | 0 | 0 | 1 | 0 | 0 | 0 |

**Table S2**: **Datasets and tasks used for pretraining T-SCAPE**

| Data Source | Dataset | Task Name | Task Explanation |
|---|---|---|---|
| IEDB | MHC I eluted ligand MS dataset | Task-EL-I | Peptide eluted from MHC masked sequence prediction |
| | MHC II eluted ligand MS dataset | Task-EL-II | |
| | MHC I peptide binding affinity dataset | Task-MHC-I | Peptide-MHC binding classification |
| | MHC II peptide binding affinity dataset | Task-MHC-II | |
| VDJdb+IEDB +McPAS-TCR +ImmuneCODE +TBAdb+10XGenomics | TCR-Peptide-MHC specificity dataset | Task-TCR | TCR-peptide-MHC binding classification |
| OAS + UNIPROT | Human non-human classification dataset | Task-Hu-MLM | Selfness conditioned masked sequence prediction |
| | | Task-Hu-CLS | Peptide Sequence selfness classification |

**Table S3**: **Datasets and tasks used for fine-tuning T-SCAPE**

| Data Source | Dataset | Task Name | Task Explanation |
|---|---|---|---|
| NCI | Peptide ELISPOT dataset | TASK-IM w/o MHC | ELISPOT result classification task |
| PRIME | MHC I ELISPOT dataset | Task-IM-I | |
| IEDB+MHCBN | MHC II ELISPOT dataset | Task-IM-II | |

**Table S4**: **The dataset's distribution of positive and negative labels according to each MHC allele type**

| Dataset | MHC Allele Type | Positive Label | Negative Label | Total Number |
|---|---|---|---|---|
| Fine-tuning Set (NCI) | - | 142 | 996 | 1138 |
| Fine-tuning Set (PRIME) | A | 1199 | 3540 | 4739 |
| | B | 287 | 1362 | 1649 |
| | C | 26 | 261 | 287 |
| Fine-tuning Set (MHCBN) | DR | 2956 | 2013 | 4969 |
| Neoantigen Discovery Set | A | 92 | 253 | 345 |
| | B | 77 | 194 | 271 |
| | C | 9 | 208 | 217 |
| Infectious Disease Set | A | 924 | 393 | 1317 |
| | B | 524 | 180 | 704 |
| | C | 40 | 32 | 72 |
